# Disordered Proteins Enable Histone Chaperoning on the Nucleosome

**DOI:** 10.1101/2020.04.17.046243

**Authors:** Pétur O. Heidarsson, Davide Mercadante, Andrea Sottini, Daniel Nettels, Madeleine B. Borgia, Alessandro Borgia, Sinan Kilic, Beat Fierz, Robert B. Best, Benjamin Schuler

**Author notes:** Address correspondence to P.O.H., R.B.B., or B.S. These authors contributed equally to this work (Pétur O. Heidarsson, Davide Mercadante).

## Abstract

Proteins with highly charged disordered regions are abundant in the nucleus, where many of them interact with nucleic acids and control key processes such as transcription. The functional advantages conferred by protein disorder, however, have largely remained unclear. Here we show that disorder can facilitate a remarkable regulatory mechanism involving molecular competition. Single-molecule experiments demonstrate that the human linker histone H1 binds to the nucleosome with ultra-high affinity. However, the large-amplitude dynamics of the positively charged disordered regions of H1 persist on the nucleosome and facilitate the interaction with the highly negatively charged and disordered histone chaperone prothymosin α. Consequently, prothymosin α can efficiently invade the H1-nucleosome complex and displace H1 via competitive substitution. By integrating experiments and simulations, we establish a molecular model that rationalizes this process structurally and kinetically. Given the abundance of charged disordered regions in the nuclear proteome, this mechanism may be widespread in cellular regulation.

---

A large fraction of the human genome codes for proteins that contain substantial disordered regions or even lack any well-defined three-dimensional structure^1^. These intrinsically disordered proteins are involved in many cellular processes and mediate key interactions with other proteins or nucleic acids^2^. DNA- and RNA-binding proteins often contain disordered regions highly enriched in positively charged residues^3,4^, which are expected to facilitate electrostatic interactions with their cellular targets, the highly negatively charged nucleic acids^3^. The affinities can be remarkably high, even if no structure is formed upon binding^5,6^. Such polyelectrolyte interactions have long been known in the field of soft matter physics^7^, but their importance in biology has only recently started to be recognized^5,6,8-11^ and is thus largely unexplored.

A ubiquitous group of proteins with long disordered positively charged regions are the histones, which are responsible for packaging DNA into chromatin. Among these, the linker histones are particularly remarkable^12^: They are largely disordered and highly positively charged, with two disordered regions flanking a small folded globular domain. By binding to the linker DNA on the nucleosome (Fig. 1a), linker histones contribute to chromatin condensation and transcriptional regulation^12,13^. However, the role of protein disorder in the complex between nucleosome and linker histone and the functional consequences have remained unclear. Here, by integrating single-molecule experiments and simulations, we establish a molecular model of the linker histone-nucleosome complex and show that disorder enables an unexpected mechanism that regulates linker histone binding: The highly negatively charged and disordered human protein prothymosin α (ProTα), a histone chaperone^14-17^ that forms a high-affinity disordered complex with linker histone H1^5^, can efficiently displace H1 from the nucleosome and accelerate its dissociation. The results can be explained by a mechanism in which ProTα invades the linker histone-nucleosome complex and forms interactions with H1 mediated by the extensive disorder in both proteins, resulting in facilitated H1 dissociation. The underlying process of competitive substitution^18^ via a transient ternary complex may be widespread in IDP-mediated nuclear interactions.

**Figure 1.**
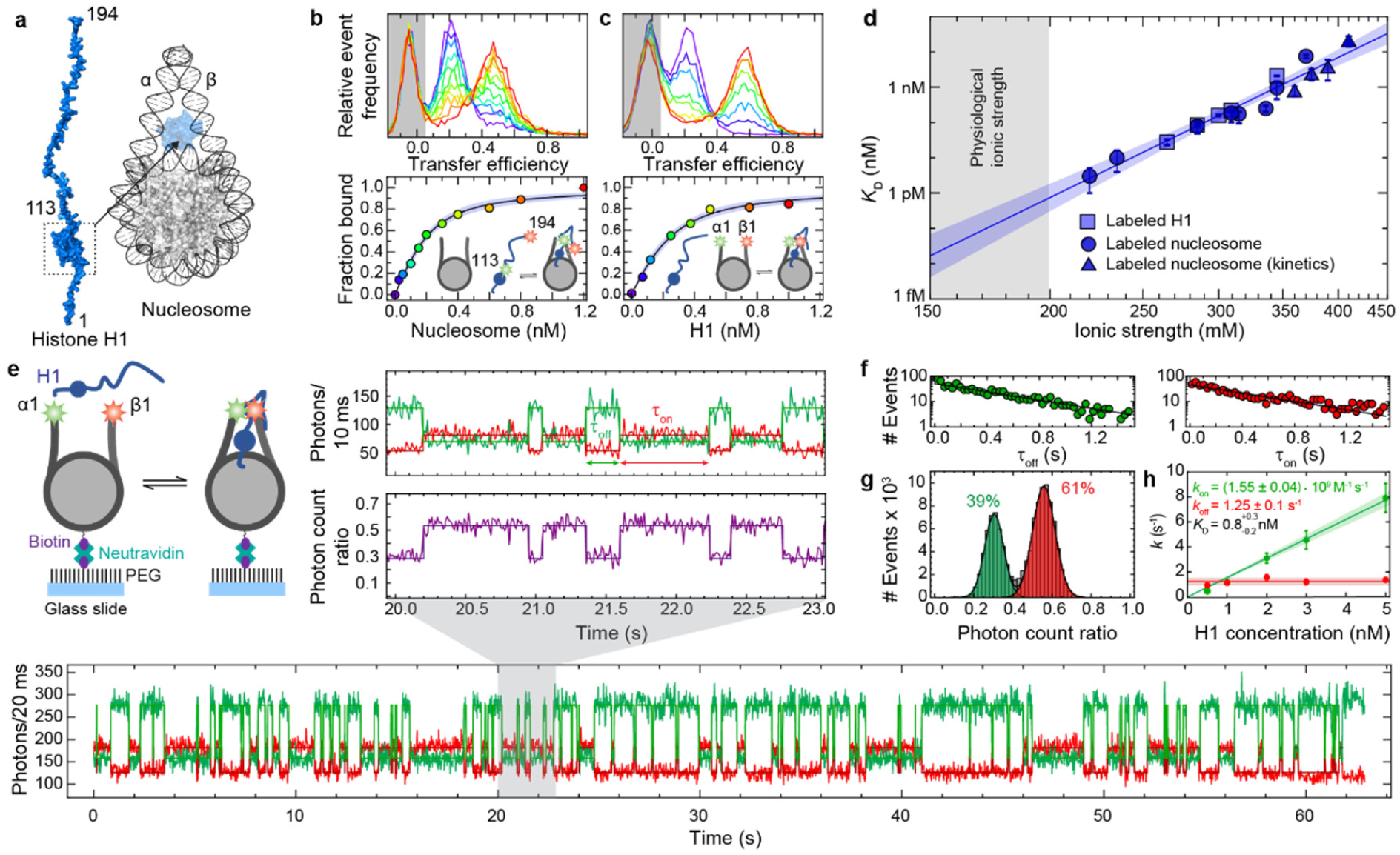
H1 binds nucleosomes tightly but reversibly. **a**, Structural representation of histone H1 and the nucleosome-H1 complex, with the position of the H1 globular domain on the nucleosome indicated (PDB 5NL0). H1: blue, DNA: dark grey, core histones: light grey. The entry and exit linker DNA are denoted α and β, respectively. **b, c**, Binding of H1 to the nucleosome. Single-molecule transfer efficiency histograms and binding isotherms (see symbols in binding isotherms for color code) of freely diffusing fluorescently labeled H1 (**b**) or labeled nucleosomes (**c**) titrated with the corresponding unlabeled ligand, at 310 mM ionic strength (10 mM Tris, 0.1 mM EDTA, pH 7.4, ionic strength adjusted with KCl). **d**, Equilibrium dissociation constant (*K*_D_) as a function of ionic strength, from binding isotherms using labeled H1 (squares) or labeled nucleosomes (circles), and from rate coefficients (*K*_D_ *= k*_off_/*k*_on_) measured with surface-immobilized labeled nucleosomes (triangles). The data were fit and extrapolated to the physiological ionic-strength range (gray-shaded area) with the Lohman-Record model,^59^ which indicates a release of 13 ± 1 counter ions upon binding (solid line). **e**, Exemplary single-molecule fluorescence time trace of 2 nM unlabeled H1 binding to surface-immobilized labeled nucleosomes at 360 mM ionic strength (acceptor signal, *N*_A_, red, donor signal, *N*_D_, green). An expanded segment is shown above, along with the photon count ratio (*N*_*A*_/(*N*_*A*_ *+ N*_*D*_), purple) and appropriately scaled state trajectories (dark green, red, purple) based on Viterbi analysis with a two-state model. **f**, Dwell-time distributions for the unbound (green) and bound states (red) with exponential fits (solid lines). **g**, Photon count ratio histograms from 46 fluorescence time traces at 360 mM ionic strength and 2 nM H1, showing the unbound (green) and bound populations (red). **h**, Association (green) and dissociation (red) rate coefficients as a function of H1 concentration at 360 mM ionic strength and fits assuming a one-to-one binding model (solid lines) with resulting values. Error bars show one standard deviation estimated from ten bootstrapping trials; shaded bands for all fits represent 95% confidence intervals.

## Probing H1-nucleosome interactions

To understand the interactions between nucleosomes, H1, and ProTα, we probed both the conformational properties of such highly disordered systems as well as the equilibria and kinetics of their complex formation and dissociation. We investigated the binding of the human linker histone H1.0 (referred to herein as H1) to nucleosomes with confocal single- molecule spectroscopy combined with Förster resonance energy transfer (FRET)^19^. We attached a donor and an acceptor fluorophore at positions 113 and 194 in H1, spanning its disordered C-terminal region (Fig. 1a), and monitored the binding to nucleosomes freely diffusing in solution.

Unbound H1 shows a low mean FRET efficiency, ⟨*E*⟩, as expected for a highly expanded configuration due to the charge repulsion within the C-terminal region (net charge +39)^5,20^. Upon addition of unlabeled reconstituted nucleosomes based on the 197-bp 601 Widom sequence^21^ (Fig. 1a), a population with higher ⟨*E*⟩ is formed (Fig. 1b), indicating a compaction of H1 on binding due to charge screening by the nucleosomal DNA, in line with previous results^22^. From the bound fraction as a function of nucleosome concentration, we obtained the equilibrium dissociation constant (*K*_D_). As expected from the electrostatic nature of the interaction, the affinity is highly dependent on ionic strength: Across the range where reliable measurements were feasible (220-375 mM), *K*_D_ increases from ∼3 pM to ∼10 nM (Fig. 1d and Supporting Information Table 1). Affinities at lower ionic strength are difficult to probe directly due to the high and nonspecific affinity of H1 to surfaces^5^, but extrapolation suggests sub-picomolar *K*_D_ values in the physiological ionic-strength range below 200 mM – even slightly tighter than estimated previously from fluorescence-based ensemble assays^23^. To control for the possible influence of dye labeling on the interaction, we also performed single-molecule measurements with the fluorophores attached at the termini of the nucleosomal linker DNA arms (positions α1 and β1, Fig. 1a) titrated with unlabeled H1 (Fig. 1c). ⟨*E*⟩ increases upon binding, in line with the expected closure of the linker DNA arms in the H1-bound nucleosome^24-26^. The resulting *K*_D_ values agree with the measurements using labeled H1 over a wide ionic strength range (Fig. 1d), attesting to the robustness of the approach and the lack of strong perturbations by the dyes.

We can thus also employ single-molecule FRET experiments for probing the kinetic mechanism of H1-nucleosome interactions. By immobilizing labeled nucleosomes via a biotin-streptavidin tether on polyethylene glycol-passivated cover slides, we were able to record fluorescence time traces with confocal single-photon counting for up to several minutes (Fig. 1e). Nucleosomal unwrapping events were identified by persistent changes in transfer efficiency^27,28^ and excluded from the analysis (Supporting Information Fig. 1c). In the presence of low concentrations of unlabeled H1, the time traces exhibit the characteristic anti-correlated changes in donor and acceptor fluorescence emission expected from the association and dissociation of individual H1 molecules, with high ⟨*E*⟩ in the bound state, and low ⟨*E*⟩ in the unbound state (Fig. 1e). To quantify the kinetics, thousands of such H1 association and dissociation events were analyzed by likelihood maximization based on a two-state Markov model^29,30^ (Fig. 1e-h; see Methods). The resulting association (*k*_on_) and dissociation rate coefficients (*k*_off_) yielded values of *K*_D_ in accord with the free-diffusion experiments (Fig. 1d and Supporting Information Table 1), indicating that surface immobilization does not interfere with H1 binding. The increase in affinity with decreasing ionic strength (Fig. 1d) is dominated by a change in *k*_off_, whereas *k*_on_ is much less salt-dependent and remains near the value expected for diffusion-limited binding (Supporting Information Fig. 2, Supporting Information Table 1).

**Figure 2.**
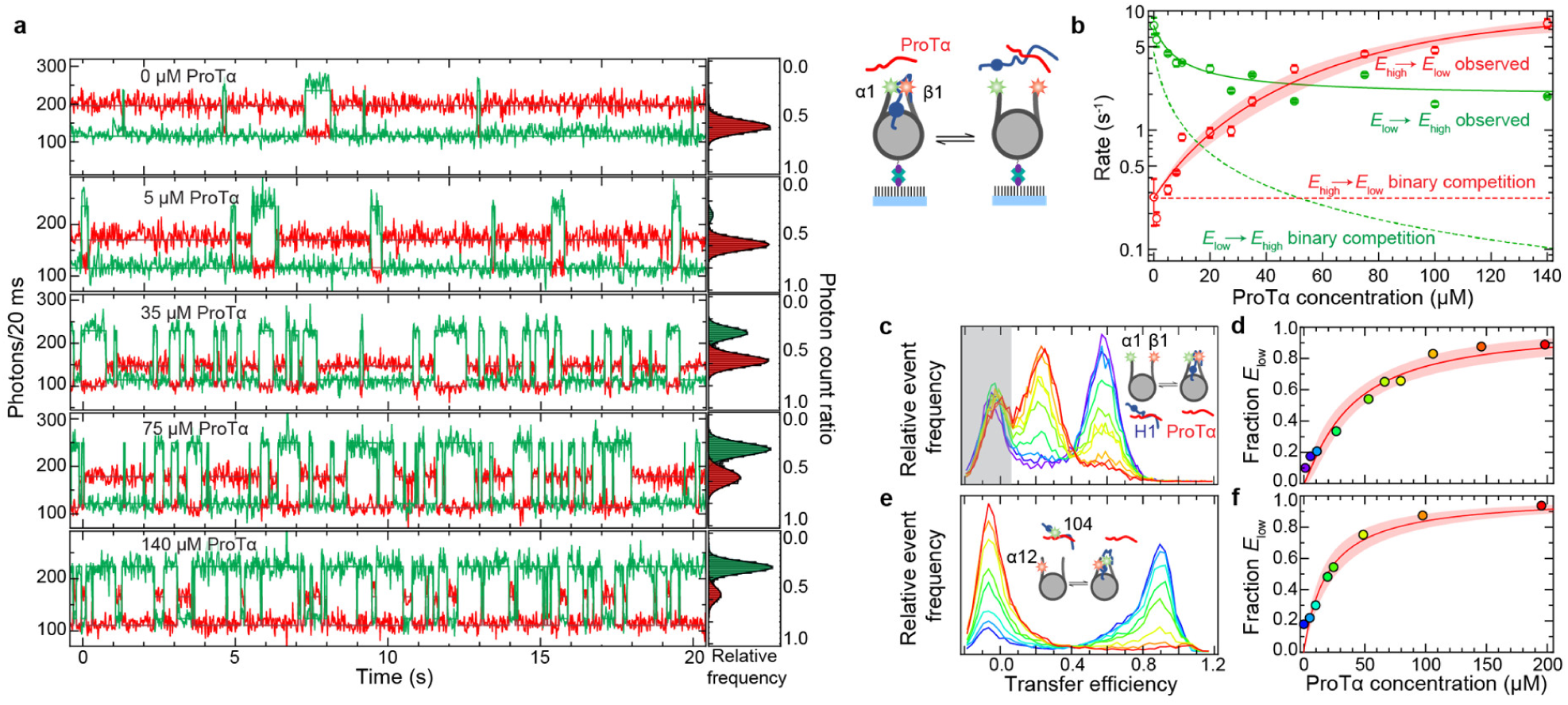
ProTα facilitates H1 dissociation from the nucleosome. **a**, Fluorescence time traces monitoring H1 binding to surface-immobilized nucleosomes (see cartoon) at different ProTα concentrations and corresponding photon count ratio histograms (*N*_*A*_/(*N*_*A*_ *+ N*_*D*_), where *N*_D_ and *N*_A_ are the numbers of photons detected in the donor and acceptor channels per 20-ms bin, respectively). **b**, Kinetic analysis of time traces, with observed transition rates from low to high FRET efficiency (*E*_low_ → *E*_high_, red symbols) and from high to low FRET efficiency (*E*_high_ → *E*_low_, green symbols) as a function of ProTα concentration, described with different kinetic models (dashed line: competition with binary interactions only; solid line: competition including ternary-complex formation; see Methods for details). Error bars show one standard deviation estimated from ten bootstrapping trials. **c-f**, Transfer efficiency histograms (c,e) from free-diffusion single-molecule FRET (see symbols in binding isotherms for color code) and corresponding fractional populations (d,f). Intramolecular FRET of double-labeled nucleosomes in the presence of 1 nM unlabeled H1, at increasing concentration of unlabeled ProTα (c) and the corresponding fraction of low-FRET population (symbols) fit with a binding isotherm (d). Intermolecular FRET between acceptor-labeled nucleosomes and donor-labeled H1, with increasing concentration of unlabeled ProTα (e) and the corresponding fraction of low-FRET population (symbols) fit with a binding isotherm (f). Note that in (c) the peak at *E* ≈ 0 (gray shading) originates from molecules lacking an active acceptor fluorophore, resulting in an amplitude independent of ProTα concentration; in (e), we explicitly use the increase in the peak at *E* ≈ 0 and concomitant decrease of the peak at *E* ≈ 0.9 to monitor H1 dissociation. Data in all panels were collected at 340 mM ionic strength, shaded bands for fits represent 95% confidence intervals.

These results illustrate a classic conundrum regarding H1-nucleosome interactions^23,31,32^: At physiological ionic strengths, dissociation of H1 from the nucleosome *in vitro* is much too slow to be compatible with efficient cellular regulation of transcription and chromatin condensation^33^. At 165 mM ionic strength, for instance, extrapolating our results yields an average dwell time in the bound state of 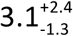 hours, similar to previous results in vitro^34^ but much longer than the timescale of about a minute observed in cells^31,32^. Clearly, other cellular factors must be involved in regulating the interaction of H1 with chromatin.

## ProTα displaces H1 from the nucleosome

A well-known linker histone chaperone is ProTα^14-17^, which binds to H1 with high affinity^5,9,35^ and is thus expected to compete with the nucleosome for H1 binding. Indeed, if the experiments probing H1-nucleosome interactions (Fig. 1e) are performed in the presence of ProTα, the kinetics of the system change markedly (Fig. 2a). For example, at 340 mM ionic strength and 3 nM H1 in the absence of ProTα, H1 remains nucleosome-bound for ∼4 s on average, and association occurs rapidly after dissociation, leading to an unbound population of only ∼2%, just enough to detect occasional dissociation events in fluorescence time traces. Upon addition of ProTα, the frequency of transitions to low ⟨*E*⟩ strongly increases (Fig. 2a), with a change in rate from 0.3 s^-1^ to 8 s^-1^ when the ProTα concentration is increased from 0 to 140 μM (Fig. 2b). This rate enhancement occurs for a wide range of ionic strengths, and is even greater at lower, near-physiological levels (Supporting Information Fig. 3a). What is the underlying competition mechanism?

**Figure 3.**
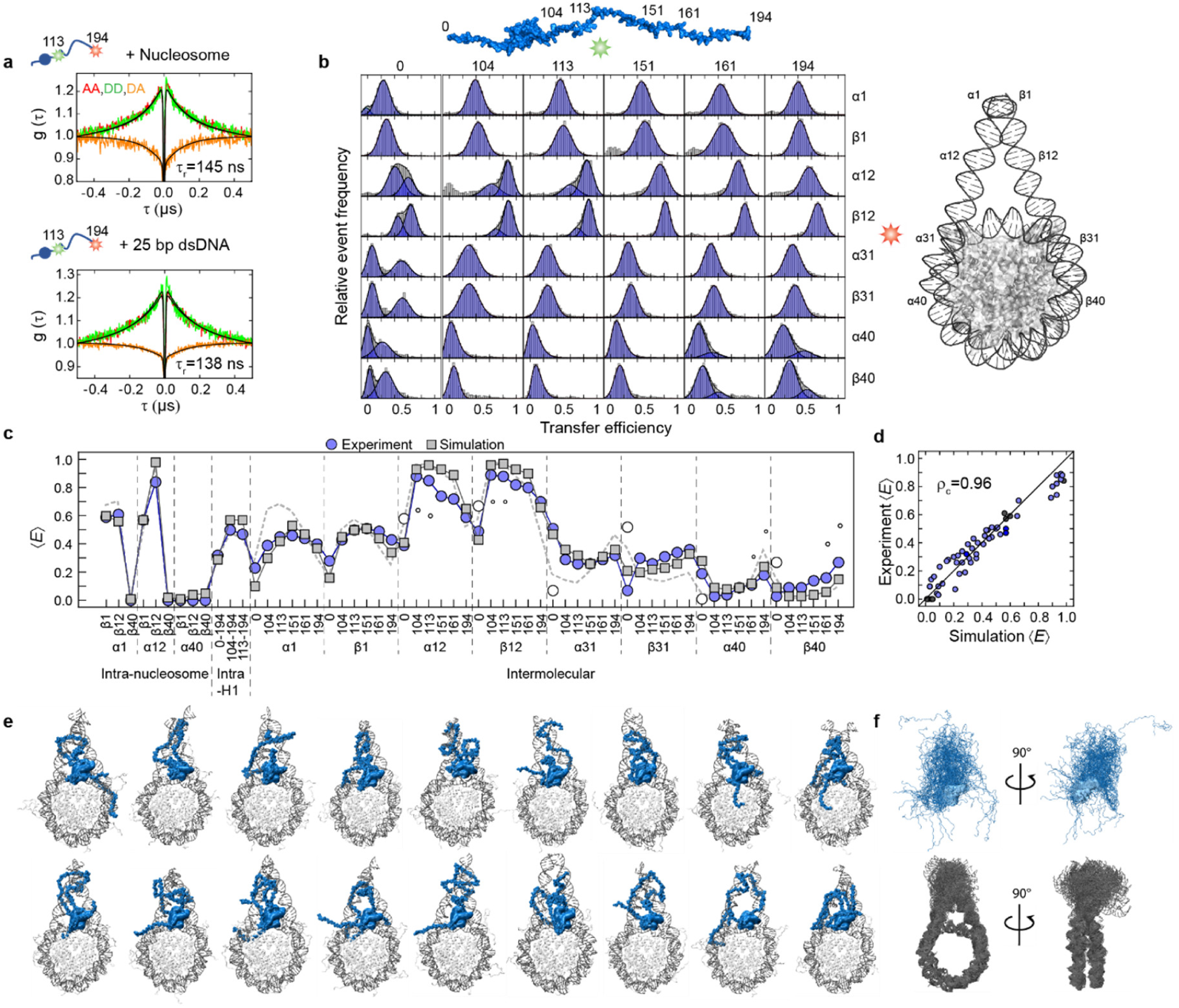
H1 remains disordered and dynamic on the nucleosome. **a**, Nanosecond fluorescence correlation spectroscopy^46,47^ of double-labeled H1 bound to the nucleosome (top) or to 25-bp double-stranded DNA (bottom), with donor-donor (DD, green), acceptor-acceptor (AA, red), and donor-acceptor (DA, orange) correlations. **b**, Transfer efficiency histograms from intermolecular single-molecule FRET experiments between donor-labeled H1 and acceptor-labeled nucleosome (labeling positions indicated along the schematic structures; numbering on the nucleosome refers to number of nucleotides from the linker DNA ends). **c**, Comparison between average FRET efficiencies, ⟨*E*⟩, from experiment and simulation for 60 different labeling pairs (b and Supporting Information Fig. 4). Filled blue circles: main experimental FRET population; Empty circles: minor experimental FRET population (size of symbol indicates population size); Gray squares: ⟨*E*⟩ from simulations using H1 globular domain orientation as in crystal structure 5NL0; Dashed line: ⟨*E*⟩ from simulations using the crystal structure but with H1 globular domain rotated by 180° on dyad (see main text and Methods for details). **d**, Correlation between experimental and simulated ⟨*E*⟩, with concordance correlation coefficient, *ρ*_c_. **e**, Representative snapshots of the H1-nucleosome complex from simulations (see Methods). **f**, Overlay of 50 conformations of H1 (upper) and nucleosome (lower) in the complex. All measurements and simulations were performed at 165 mM ionic strength.

The observed increase in rate is not compatible with a simple competition mechanism where H1 forms only binary complexes, either with the nucleosome or with ProTα. In this case, H1 would first have to dissociate from the nucleosome before binding to ProTα. In other words, dissociation of H1 from the nucleosome would be a unimolecular reaction with a rate independent of ProTα concentration (red dashed line in Fig. 2b), in contrast to what we observe. Moreover, the observed rate of transitions to high ⟨*E*⟩ changes much less than expected for the association rate in simple competition, where the presence of ProTα should sequester free H1 in solution to such an extent that H1 association decreases to a virtually undetectable level (green dashed line in Fig. 2b). These deviations in the kinetics from the simple competition mechanism are a typical indication for the formation of ternary complexes, a mechanism that has recently emerged as a paradigm for gene regulation and typically involves multivalent interactions^36-40^.

Ternary complex formation immediately suggests itself for IDPs such as H1 and ProTα, whose interactions can be considered an extreme case of multivalency^8,36,40,41^: If the positively charged tails of H1 remain largely disordered and dynamic while bound to the nucleosome, as suggested by recent work on H1 bound to double-stranded DNA^6^ and from simulations^26^, the negatively charged ProTα would still be able to invade the complex and bind to H1 while remaining disordered^5^. Intra- and intermolecular single-molecule FRET experiments show that ProTα binding indeed accelerates dissociation of H1 from the nucleosome (Fig. 2c, Supporting Information Fig. 3). Notably, the ProTα concentrations required for enhanced H1 dissociation are in the physiological range^42^ and are thus likely to contribute to the high mobility of H1 in cells^31,32^. To obtain more insight into the underlying molecular mechanism of ProTα action, we first require a structural description of H1 on the nucleosome.

## Highly disordered H1 on the nucleosome

The globular domain of H1 is known to localize to the dyad axis and interact with the nucleosomal core DNA and both linkers^24,25,43^. However, the conformational distributions of the disordered regions of H1 on the nucleosome have been more difficult to elucidate because of their pronounced dynamics and the lack of persistent structure^4,6,25,44,45^. We can monitor the presence of dynamics in single-molecule FRET experiments with nanosecond fluorescence correlation spectroscopy (nsFCS)^46^, since fluctuations in distance cause fluctuations in the fluorescence intensity of donor and acceptor^47^. Indeed, H1 shows pronounced long-range chain dynamics on the 100-ns timescale characteristic of disordered proteins^47^, both in isolation^5^, bound to the nucleosome (Fig. 3a), and bound to a 25-bp double-stranded DNA (in agreement with both recent NMR results^6^ and the insensitivity of intermolecular DNA/H1 FRET efficiency to label position, Supporting Information Fig. 4). Dynamics on this timescale can be observed throughout the C-terminal region of H1, as well as between the ends of the nucleosomal linker DNA (Supporting Information Fig. 4), highlighting the exceedingly dynamic nature of the complex. The presence of broad distance distributions and rapid dynamics resulting from disorder is further supported by fluorescence lifetime analysis (Supporting Information Fig. 5).

**Figure 4.**
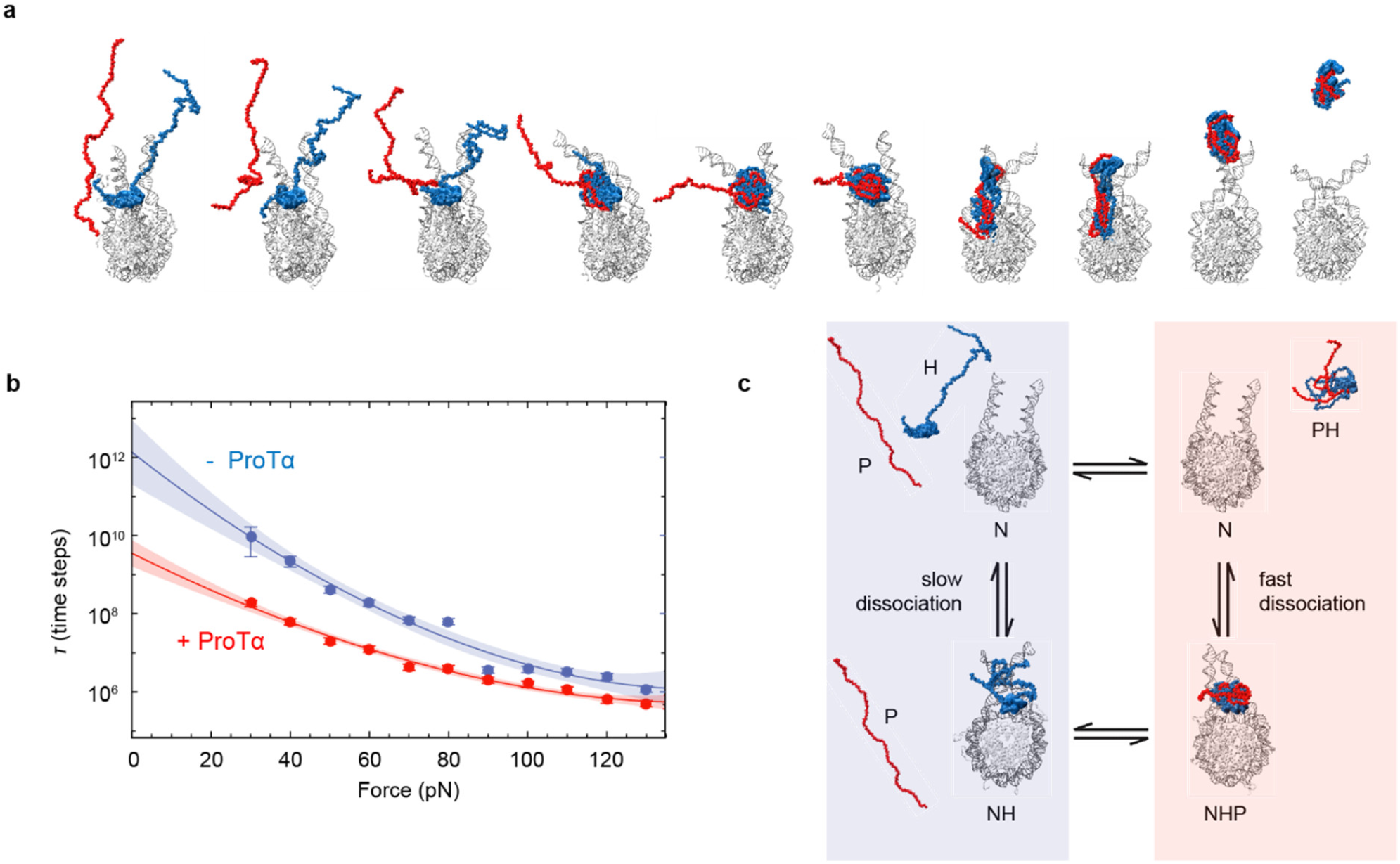
Mechanism of H1 chaperoning on the nucleosome by ProTα. **a**, Structural model based on simulation snapshots depicting the association of ProTα (red) to the H1-nucleosome complex followed by the dissociation of ProTα-H1 from the nucleosome. **b**, Escape time of H1 from the nucleosomes in simulations as a function of pulling force in the absence (blue) and presence (red) of ProTα (see Methods for details). Dissociation in **a** was at 40 pN. **c**, Kinetic scheme showing the relevant molecular states. H1 (H) binds to the nucleosome (N) rapidly, leading to condensation of the C-terminal tail of H1 and a collapse of the linker DNA. H1 dissociates extremely slowly from the nucleosome by itself, but binding of ProTα (P) to H1 on the nucleosome facilitates dissociation of H1 in the ProTα-H1 complex (PH).

We thus require a molecular model that takes these dynamics into account, allows a description of H1 in terms of a structurally diverse conformational ensemble, and can be optimized based on experimental data. In view of the large size of the system, we employed a coarse-grained model that captures both the known structure of the nucleosome and the pronounced dynamics of the linker DNA and the disordered regions of H1. We used a “top-down” coarse-grained model based on matching experimental observations, rather than a “bottom-up” coarse-grained model matched to all-atom simulations^26^. We combined a structure-based model^48,49^ for the nucleosomal core particle and the globular domain of H1 and its position on the dyad based on the crystal structure (PDB: 5NL0)^25^ with a polymer-like representation of the histone tails, which was previously used successfully to describe the interaction of H1 with ProTα^5^. The interaction between disordered regions of histones and DNA was encoded in terms of nonspecific short-range and electrostatic interactions^50^, including a screening term to account for the experimental ionic strength (see Methods). To attain a realistic description of the disordered and dynamic parts of the H1-nucleosome complex, we adjusted the strength of the short-range protein-DNA interactions via a single parameter to maximize the agreement between the measured FRET efficiencies and those computed from the simulation ensemble.

To test and optimize the model of H1 on the nucleosome, we obtained a total of 60 intra- and intermolecular distance constraints from single-molecule FRET experiments. We created constructs with labeling sites on the nucleosomal DNA, both in positions corresponding to the linker DNA and in the core particle, and along the H1 sequence (Fig. 3b, Supporting Information Fig. 4, Supporting Information Table 2). Intermolecular FRET experiments using alternating excitation of donor and acceptor^51^ (see Methods) indicate 1:1 H1-nucleosome complexes under our experimental conditions (Supporting Information Fig. 5), in accord with previous results^23^. Most of the resulting transfer efficiency histograms exhibit a single peak, as expected from the rapid conformational averaging within the complex observed by nsFCS (Fig. 3a)^46^. The agreement between transfer efficiencies involving rotationally symmetric positions of the labels on the α and β linker DNA attests to the robustness of the results.

The conformational ensembles resulting from the optimized simulation model show very good agreement with the measured FRET efficiencies and other experimental data. In the H1-bound state, the characteristic pattern of transfer efficiencies for different labeling positions along the H1 and DNA sequences are described remarkably well by this simple model (Fig. 3c and Supporting Information Fig. 6), and even the absolute efficiency values show excellent overall agreement (concordance correlation coefficient 0.96, Fig. 3d). The simulations also reproduce the increase in FRET efficiency between the labeled linker DNA arms observed experimentally upon binding of H1 (Fig. 3c and Supporting Information Fig. 4e), in accord with electron cryo-microscopy results that show a compaction of the nucleosome and crossing of the linker DNA arms when H1 is bound^25^.

In some cases, reproducible broadening or double peaks were observed in the transfer efficiency histograms (Fig. 3b), especially if the labels are located close to the exit and entry points of the nucleosomal DNA and to the globular domain of H1. We hypothesized that they are caused by the different orientations of the H1 globular domain on the dyad that were previously identified by crosslinking experiments and are expected to interconvert very slowly^12,13,25^. Indeed, when the simulations were performed with the globular domain of H1 rotated by 180° on the dyad compared to the crystal structure^25^, the FRET efficiencies of several of the secondary populations could be well reproduced (Fig. 3c). Explicitly including a representation of the fluorophores in the simulations yielded very similar results (Supporting Information Fig. 6). Overall, these results indicate that this coarse-grained, charge-dominated description of the disordered regions of H1 on the nucleosome captures the essential properties of the ensemble and further support the notion that H1 retains its disorder when bound to the nucleosome^6,25,52^. Figure 3e illustrates the broad range of conformations that are populated in the H1-nucleosome complex.

## Disorder enables histone chaperoning

The disorder and the large-amplitude conformational fluctuations of the positively charged regions of H1 on the nucleosome suggest a simple mechanism for the interactions with its negatively charged chaperone ProTα. A possible sequence of events can be obtained by including ProTα in the simulations (Fig. 4): Once the initial contacts are formed in a fly-casting-like process^53^, ProTα increasingly interacts with H1. As a result, it competes with the electrostatic interactions between the linker DNA and the disordered regions of H1, thus effectively reducing the interaction strength of H1 with the nucleosome and leading to an opening of the linker DNA arms. Since spontaneous dissociation of H1 was too slow to be observed on the timescale of the simulations, we accelerated dissociation by applying a pulling force between the nucleosome and the globular domain of H1 (see Methods) and compared the dissociation times of H1 as a function of applied force with and without ProTα (Fig. 4). H1 dissociation is consistently faster in the presence of ProTα than in its absence (Fig. 4 and Supporting Information Fig. 6), in agreement with experiment.

Our results suggest that disordered proteins facilitate a remarkable mechanism of molecular competition: highly disordered, electrostatically driven biomolecular complexes aid the formation of transient ternary interactions, where a competing binding partner invades an existing complex and accelerates dissociation. This mechanism of ‘competitive substitution’ has been known in the field of synthetic polyelectrolytes^18^, but its role in biology has largely remained unexplored. Given the great abundance of disorder in nuclear proteins, including transcriptional regulators, polymerases, and other RNA- and DNA-binding factors^4^, competitive substitution is likely to be widespread among biomolecular interactions in the nucleus and may play an important role in cellular regulation. Even though the existence of charged disordered regions has been known for decades, we are only now starting to elucidate the physical basis underlying their functions. The physical properties of the charged disordered regions of H1 and the core histones are also likely to be crucial for processes involving liquid-liquid phase separation in chromatin^8,10,11,54,55^.

How relevant is the effect of ProTα in a cellular environment? Clearly, a multitude of contributions are likely to modulate the interaction of H1 with chromatin, such as posttranslational modifications^56^, other histone chaperones^16^, and DNA-interacting machinery^4^, or local variations in ion concentrations^57^. However, several lines of evidence indicate that ProTα is an important effector in vivo. First, ProTα has been shown to increase H1 mobility in cells^17^; second, the range of concentrations where we observe ProTα to change H1-nucleosome affinity correspond to the reported intracellular concentrations^42^; and finally, the facilitated H1 dissociation we observe is a large effect and robust to salt concentration (Supporting Information Fig. 3). Correspondingly, variations in ProTα concentration, e.g. during the cell cycle^58^, may be involved in regulating H1 association with chromatin. Irrespective of the absolute ProTα concentration, another observation is noteworthy: the affinity of H1 to free DNA is about two orders of magnitude lower (Supporting Information Fig. 4) than its affinity to nucleosomes (Fig. 1). As a result, ProTα is expected to efficiently prevent nonspecific binding of H1 to DNA and ensure its targeting to the nucleosomal dyad.

Finally, our results suggest that two long-standing questions in the chromatin field, namely the nature of the structural ensemble of H1 on the nucleosome^8^ and the discrepancy between the residence times of H1 in vivo^31^ and in vitro^33^, are closely connected: It may be precisely the large degree of structural disorder and long-range dynamics in the tails of nucleosome-bound H1 that enable chaperones such as ProTα to invade the H1/nucleosome complex and in this way accelerate H1 dissociation by competitive substitution instead of passively scavenging H1 once dissociated. Related processes involving charged disordered proteins may affect many aspects of chromatin assembly and dynamics, and cellular regulation in general^10,11,26,55^.

## SUPPLEMENTARY INFORMATION

is available in the online version of the paper.

## ACKNOWLEDGEMENTS

We thank Iwo König for providing ProTα samples, Karin Buholzer and Flurin Sturzenegger for helpful discussion, Fabienne Büchler and Nathalie Wyss for excellent technical assistance, and the Functional Genomics Center Zurich for performing mass spectrometry. This project was funded by the Novo Nordisk Foundation (P.O.H.), the Carlsberg Foundation (P.O.H.), The Boehringer Ingelheim Fonds (S.K.), the Swiss National Science Foundation (B.S. and B.F.), Ecole Polytechnique Fédérale de Lausanne (B.F.), and the Intramural Research Program of the NIDDK at the National Institutes of Health (R.B.B.). This work utilized the computational resources of the NIH HPC Biowulf cluster (http://hpc.nih.gov) and of Piz Daint at the CSCS Swiss National Supercomputing Centre.

## AUTHOR CONTRIBUTIONS

P.O.H., D.M., R.B.B, and B.S. designed research; P.O.H and S.K. prepared reconstituted nucleosomes; P.O.H., M.B., A.B., A.S., S.K., and B.F. prepared fluorescently labeled and/or unlabeled proteins; P.O.H. and A.S. performed single-molecule experiments; P.O.H., A.S., D.N., and B.S. analyzed single-molecule data; D.M. and R.B.B. performed and analyzed simulations; R.B.B, B.F., and B.S. supervised research; P.O.H. and B.S. wrote the paper with help from all authors.

## Methods

### Protein preparation and labeling

Recombinant wild-type human histone H1.0 (New England Biolabs, cat. # M2501S) was used for experiments with fluorescently-labeled nucleosomes. ProTα was prepared as previously described^20^. Variants of H1 for fluorescent labeling were produced by bacterial expression using a modified version of the pRSET vector^60^ containing the human H1F0 gene (UniProt P07305), a hexahistidine tag and a thrombin cleavage site. All protein variants were confirmed to have the correct molecular weight by mass spectrometry. Cysteine mutations were introduced for labeling the protein with fluorescent dyes using site-directed mutagenesis. H1 variants were expressed, purified, and fluorescently labeled as previously described^5^. Briefly, all H1 variants were expressed in in *Escherichia coli* C41 cells in terrific broth medium at 37 °C, using isopropylthiogalactopyranoside (IPTG) to induce expression. The cells were pelleted after three hours of growth, resuspended in denaturing buffer containing 6 M guanidinium chloride (GdmCl) and the soluble fraction applied to a Ni-IDA resin (Agarose Bead Technologies). After washing, the protein was eluted using 250-500 mM imidazole, then dialyzed against phosphate-buffered saline (PBS), followed by thrombin (Serva) cleavage to remove the hexahistidine tag. Uncleaved protein and cleaved hexahistidine tag were removed from the eluted mixture with a HisTrap HP 5 mL column (GE Healthcare) in PBS with 25 mM imidazole. The final purification step was performed by anion exchange chromatography using a Mono S column (GE Healthcare). Before fluorescent labeling, samples were reduced with dithiothreitol (DTT) and purified by reversed-phase high-performance liquid chromatography (RP-HPLC) using a Reprosil-Gold C4 column. H1-containing fractions were resuspended and labeled in denaturing buffer (6 M GdmCl, 50 mM sodium phosphate, pH 7.0), before a final RP-HPLC purification step. Lyophilized proteins were resuspended in 8 M GdmCl, frozen in liquid nitrogen and stored at −80 °C. All experiments were performed in TEK buffer (10 mM Tris, 0.1 mM EDTA, pH 7.4) at different ionic strengths adjusted by addition of KCl. Protein sequences and labeling positions are shown in Supporting Information Table 2. The dye pairs Alexa Fluor 488/Alexa Fluor 594 (Förster radius *R*0 = 5.4 nm) and Cy3B/CF660R (*R*0 = 6.0 nm) were used for experiments with freely-diffusing molecules and for surface-immobilized nucleosomes, respectively.

### Preparation of fluorescently labeled oligonucleotides

A ∼10-µL solution of 5-10 nmol of oligonucleotide (thymine modified with a C6-amino linker for the reaction with the succinimidyl ester of the fluorescent dye, Integrated DNA Technologies, see Supporting Information Table 2) was diluted with 50 µL of labeling buffer (0.1 M sodium bicarbonate, pH 8.3). For analytical purposes, a 1-µL sample was diluted with 50 µL of RP-HPLC solvent A (95% triethylammonium acetate/5% acetonitrile), followed by RP-HPLC using a Reprosil-Pur 200 5 µm, 250 x 4mm C18 column (Dr. Maisch) using a gradient of 0-100% RP-HPLC solvent B (100% acetonitrile) in 50 minutes. 50-100 µg of fluorescent dye succinimidyl ester dissolved in dimethylsulfoxide were sonicated for 10 minutes and added to the oligonucleotide solution in labeling buffer and the reaction incubated at room temperature for at least two hours. After ethanol precipitation to remove excess dye, the pelleted oligonucleotide was redissolved in 100 µL 95%A/5%B RP-HPLC solvent. Labeled oligonucleotides were purified using the same column and gradient as above, lyophilized, and resuspended in double-distilled water (ddH_2_O) to a final concentration of 2.5 µM and stored at −20 °C. The correct molecular weights of the labeled oligonucleotides were confirmed by mass spectrometry.

### Preparation of core-histone octamer

Human wild-type core histones (H2A, H2B, H3, H4) were prepared as previously described^61^. Each core histone protein was expressed in BL21 De3 pLysS cells using a pET3a plasmid carrying the corresponding gene. Cells were grown at 37 °C in LB media including 100 µg/mL ampicillin and 35 µg/mL chloramphenicol until an OD600 of 0.6 was reached, at which point protein expression was induced by addition of IPTG to a final concentration of 0.5 mM. The cells were harvested 3 hours after induction, the cell pellets resuspended in lysis buffer (20 mM Tris, 1 mM EDTA, 200 mM NaCl, 1 mM 2-mercaptoethanol (2−ME), one cOmplete protease inhibitor cocktail tablet (Roche) per 50 mL, pH 7.5), followed by cell lysis by freeze-thawing and sonication. The inclusion body pellet was washed three times with the histone lysis buffer (twice with, and once without 1% triton X-100). The histones were resolubilized in histone resolubilization buffer (6M GdmCl, 20 mM Tris, 1 mM EDTA, 1 mM 2−ME, pH 7.5) and then dialyzed against urea buffer (7 M urea, 10 mM Tris, 1 mM EDTA, 0.1 M NaCl, 5 mM 2−ME, pH 7.5). The proteins were then purified by cation exchange using a 5-mL HiTrap SP HP column (GE Healthcare) and the collected fractions analyzed by SDS-PAGE. The final purification step was performed by preparative RP-HPLC on a Zorbax 300SB 7μm 21.2×250mm column using a gradient of 30-70% solvent B (A: water with 0.1% TFA, B: 9.9% water, 90% acetonitrile with 0.1% TFA). The collected fractions were characterized by analytical RP-HPLC and ESI-MS, lyophilized, and stored at −20 °C.

To refold the core-histone octamer, 0.4-1.5 mg of each of the purified lyophilized human histones were dissolved in 6 M GdmCl, 10 mM Tris, 5 mM DTT, pH 7.5, and the protein concentrations quantified by UV absorbance. Equimolar amounts of H3 and H4 were mixed with 1.05 equivalents of H2A and H2B to a final concentration of 1 mg/mL, and octamers were refolded by dialyzing against 2 M NaCl, 10 mM Tris, 1 mM EDTA, 5 mM DTT, pH 7.5. The refolded octamers were then purified by gel filtration on a Superdex S200 10/300GL column (GE Healthcare) and the collected fractions analyzed by SDS-PAGE. The octamer-containing fractions were pooled and concentrated to ∼50 µM; glycerol was added to a final concentration of 50% (vol/vol) and the samples stored at −20 °C.

### PCR amplification of 197-bp DNA containing the 601 Widom sequence

DNA for nucleosome reconstitution was generated by PCR amplification of a pJ201 plasmid template containing the 147 bp Widom sequence^21^, either with unlabeled (Integrated DNA Technologies, USA) or fluorescently-labeled oligonucleotides (see *Preparation of fluorescently labeled oligonucleotides*). DNA for nucleosomes for surface-immobilization was generated by using oligonucleotides with a modified thymidine containing a C6-aminolinker and a biotin moiety (Integrated DNA Technologies). The oligonucleotides were designed so that the Widom sequence was extended by linker DNA of 25 bp length on either side. The PCR was typically performed in 10 x 50 µL volume by mixing in PCR tubes Phusion HF buffer (1x, New England Biolabs), plasmid template (0.02 ng/µL), 0.25 µM forward primer, 0.25 µM reverse primer, and dNTPs (0.2 mM each) with ddH_2_O and 1.0 units of Phusion high-fidelity DNA polymerase (New England Biolabs). Thermocycles included 30 s for initial denaturation at 94 °C, followed by 30 cycles of 20 s for denaturation at 94 °C, 10 s for annealing at 66 °C, and 15 s for extension at 72 °C. The completed PCR reactions were pooled and ethanol-precipitated prior to purification using a DNA Clean & Concentrator Kit (DCC-25, Zymo Research). The concentration of the labeled 197-bp PCR products was determined by UV absorbance. DNA sequences of the 197-nucleotide α and β strands containing the 601 Widom sequence, primer sequences, and positions of nucleotides that were modified for labeling or biotinylation are listed in Supporting Information Table 2.

### Nucleosome reconstitution

Nucleosomes were reconstituted^62^ using 10 pmol of purified DNA containing the 147-bp 601 Widom sequence^21^ flanked by 25-bp linkers. The DNA was mixed with 0.9-1.5 molar equivalents of recombinant core histone octamer at a final concentration of 2 M NaCl on ice. The 30-µL reaction was then transferred to a Slide-A-Lyzer MINI dialysis device (Thermo Fisher Scientific) and dialyzed against a linear gradient of buffer with decreasing salt concentration, starting from 10 mM Tris, 0.1 mM EDTA, 2 M KCl, pH 7.5 to 10 mM Tris, 0.1 mM EDTA, 10 mM KCl, pH 7.5 over ∼20 hours. The gradient was created by slowly removing buffer from the dialysis container with constant flow rate using a peristaltic pump and, simultaneously, supplying fresh buffer with 10 mM KCl using the same flow rate and thus keeping the volume constant. The reactions were then transferred to microcentrifuge tubes and centrifuged for 5 mins at 21000 x g and 4 °C to remove aggregates; the supernatant was transferred to a new tube. After determining the volumes and concentrations of the samples via absorbance at 260 nm, 0.2-0.5 pmol of the reaction products were loaded on a 6% agarose gel (Invitrogen) and run for 90 mins at 90 V with 0.25x Tris-Borate as running buffer. The gels were stained with GelRed (Biotium) for 30 minutes and visualized under UV light. Only nucleosome preparations containing less than 5% of free DNA in the sample were used for measurements.

### Surface immobilization of nucleosomes

Quartz coverslips coated with polyethylene glycol and biotin (MicroSurfaces) were sonicated in Tween-20 (Thermo Fisher Scientific) and extensively washed with ddH_2_O before binding to silicone hybridization chambers (SecureSeal™ hybridization chambers, Grace BioLabs), forming chambers with a sample volume of 150 µL. The chambers were washed several times with 50 mM sodium phosphate buffer containing 0.01% Tween-20 and then incubated with a 1-µM neutravidin solution (Vector Labs) for 10 minutes. After three washing steps with TEK buffer including 0.01% Tween-20, the chambers were filled with a 10- to 20-pM solution of biotinylated and fluorescently labeled nucleosomes in TEK buffer with 0.1 mg/mL bovine serum albumin (BSA) and 0.01% Tween-20 for 5-10 minutes, followed by another two washing steps in TEK buffer.

### Single-molecule fluorescence spectroscopy

Single-molecule experiments on freely diffusing molecules were conducted at 22 °C using either a custom-built confocal instrument or a MicroTime 200 (PicoQuant), including a HydraHarp 400 time-correlated single photon counting module (PicoQuant). The donor dye was excited using a 485-nm diode laser at 100 µW power (measured at the back aperture of the objective), either in continuous-wave mode or with pulsed interleaved excitation^51^ to enable alternating excitation of donor and acceptor dyes. For acceptor excitation, the light from a supercontinuum laser (NKT Photonics) operating at 20 MHz repetition rate was passed through a z582/15 band-pass filter (Chroma) and adjusted to an average power of 35 µW at the back aperture of the objective. Excitation and emission light was focused and collected, respectively, using a high-numerical-aperture microscope objective (Olympus UplanApo 60x/1.20 W). Emitted fluorescence was focused onto a 100-µm pinhole and separated into four channels by polarization and donor and acceptor emission wavelengths. Single-photon avalanche diodes were used for detection (SPCM-AQR-15, PerkinElmer, or τ-SPADs, PicoQuant). All experiments involving freely diffusing molecules were performed in µ-slide sample chambers (ibidi) at 22 °C in 10 mM Tris, 0.1 mM EDTA, pH 7.4 with varying ionic strength by adjusting the KCl concentration to 150-400 mM; 140 mM 2−ME and 0.01% (vol/vol) Tween-20 were added for photoprotection and for minimizing surface adhesion, respectively.

Single-molecule experiments with surface-immobilized nucleosomes were conducted on a custom-built confocal instrument with a 532-nm continuous-wave laser (LaserBoxx LBX-532-50-COL-PP, Oxxius). The objective (UPlanApo 60x/1.20-W, Olympus) was mounted on a piezo stage (P-733.2 and PIFOC, Physik Instrumente GmbH) for scanning. Fluorescence emission was collected and split into two channels with a dichroic mirror (T635LPXR, Chroma). Donor emission was filtered with an ET585-65m bandpass filter (Chroma) and detected with a τ-SPAD (PicoQuant); acceptor emission was filtered with a LP647RU long-pass filter (Chroma) and detected with a SPCM-AQRH-14 single-photon avalanche diode (Perkin Elmer). After surface immobilization of nucleosomes, experiments were performed by filling the reaction chamber with degassed TEK buffer containing H1 and/or ProTα, under argon atmosphere to exclude atmospheric oxygen. The buffer contained KCl to adjust the ionic strength, 0.1 mg/mL of BSA to reduce nonspecific surface interactions, and D_2_O was used instead of H_2_O to increase the quantum yield of the dyes^63^. An oxygen scavenging system containing 2.5 mM protocatechuic acid and 1 nM protocatechuate-3,4-dioxygenase was used to improve photostability^64^, and 1 mM methyl viologen and 1 mM ascorbic acid^65^ were used for triplet quenching and radical scavenging. A surface area of 20 x 20 µM was scanned (78 nm/pixel) at a laser power of 2 µW to locate single immobilized nucleosomes. Z-position and objective correction collar were adjusted to maximize the fluorescence intensity detected.

Data for FRET efficiency histograms from freely diffusing molecules were collected on samples containing 50-100 pM double-labeled H1 or nucleosomes; inter-molecular FRET efficiencies were measured on samples with up to 500 pM of acceptor-labeled nucleosomes to ensure saturation of binding, and up to 5 nM in time-resolved FRET experiments at elevated ionic strengths (Supporting Information Fig. 3). FRET efficiencies were calculated according to 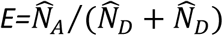, where 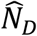 and 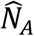 are the number of donor and acceptor photons in the fluorescence burst, respectively, after correction for background, direct acceptor excitation, channel crosstalk, differences in dye quantum yields, and different efficiencies of photon detection^66^. Aggregates, identified as occasional fluorescence bursts with photon counts greater than three standard deviations from the mean signal binned at 1s, were removed before data analysis. For experiments on surface-immobilized nucleosomes, we determined the photon count ratio, summed for all traces recorded at the same ionic strength and H1 concentration, which reports accurately on the ratio of bound and unbound populations (even though the photon count ratio may differ from the corrected absolute transfer efficiencies).

### Nanosecond fluorescence correlation spectroscopy (nsFCS)

Data for nsFCS^46,47^ were collected using continuous-wave excitation at 485 nm and a ∼100-pM sample of fluorescently labeled H1, either with acceptor-labeled nucleosomes or with an excess of unlabeled dsDNA or nucleosomes. Donor and acceptor fluorescence photons from the bound subpopulation were used for the correlations at 1 ns binning time. Photons were recorded with two detectors each for donor and acceptor and cross-correlated between detectors to avoid the effects of detector dead times and after-pulsing on the correlation functions. Autocorrelation curves of acceptor and donor channels and cross-correlation curves between acceptor and donor channels were fit and analyzed as described previously^5^. Briefly, the correlation curves were fit over a time window of 2.5 µs using

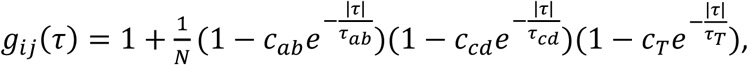

where *i* and *j* indicate donor (*D*) or acceptor (*A*) fluorescence emission; *N* is the effective mean number of molecules in the confocal volume; *c*_*ab*_, *τ*_*ab*_, *c*_*cd*_ and *τ*_*cd*_ are the amplitudes and time constants of photon antibunching (ab) and chain dynamics (cd), respectively; *c*_*T*_ and *τ*_*T*_ refer to the triplet blinking component occurring on the microsecond timescale. Fluctuations in the interdye distance result in specific features in the correlation functions: donor and acceptor autocorrelations have a positive amplitude (*c*_*cd*_ > 0), whereas the donor-acceptor cross-correlation has a negative amplitude (*c*_*cd*_ < 0), but all correlations have identical decay times^67^. All three correlation curves were thus fit globally with the same value of *τ*_*cd*_, but with *c*_*cd*_, *c*_*ab*_, *τ*_*ab*_, *τ*_*T*_, and *c*_*T*_ as free fit parameters. *τ*_*cd*_ can be converted to the reconfiguration time of the chain, *τ*_*r*_, by assuming that the chain dynamics can be modeled as a diffusive process in the potential of mean force derived from the sampled inter-dye distance distribution *P*(*r*)^46,67^ based on the SAW-v model^68,69^. The correlation functions were normalized to 1 at their respective values at 0.5 μs to facilitate direct comparison.

### Fluorescence lifetime and fluorescence anisotropy

Fluorescence lifetimes were estimated from the mean donor (⟨*t*_*D*_⟩) and acceptor (⟨*t*_*A*_⟩) arrival times after the excitation pulse. The average lifetimes were then combined with the transfer efficiencies in two-dimensional plots (Supporting Information Fig. 5) where 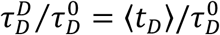 and 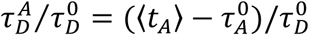 were calculated for each burst. Here, 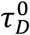 and 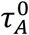 are the intrinsic donor and acceptor lifetimes, respectively. For a fixed interdye distance, the normalized mean fluorescence lifetimes are equal to 1-*E* (diagonal lines in Supporting Information Fig. 5), whereas systems rapidly sampling a broad distance distribution deviate from the diagonal^*70,71*^.

Fluorescence anisotropies for donor and acceptor dyes were determined for all intra- and inter-molecular labeling pairs from the single-molecule measurements and yielded values of 0.09-0.14 for the donor dye and 0.16-0.22 for the acceptor dye (upon direct excitation), which indicates sufficiently rapid reorientational dynamics of the dyes for approximating the orientational factor in Förster theory by 2/3^72^.

### Binding affinities

Transfer efficiency histograms were recorded for either doubly labeled H1 or nucleosomes with increasing concentrations of the unlabeled binding partner until no further change in transfer efficiency was observed. The histograms were fit with two or more Gaussian peak functions to quantify the areas of the bound and unbound subpopulations and the resulting fraction of bound species (*θ*). The binding isotherm (*θ* vs. ligand concentration) was then fit to quantify the dissociation constant (*K*_D_) using

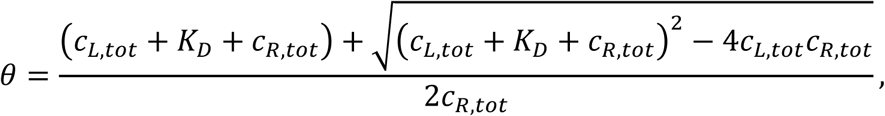

where *c*_L,tot_ and *c*_R,tot_ are the total concentrations of H1 or nucleosome, depending on which molecule is kept at constant concentration (*c*_L,tot_ variable, *c*_R,tot_ constant). Errors on *K*_D_ values are based on pipetting error estimates, propagated through corresponding dilution steps at each ionic strength. The number of counterions released upon complex formation can be estimated from the ionic-strength dependence of the *K*_D_ according to the model of Record et al.^59^.

### Analysis of fluorescence time traces from surface-immobilized molecules

Single-molecule fluorescence time traces were first inspected visually, and any recordings showing pronounced changes in count rate, i.e. those indicative of nucleosome unwrapping (Supporting Information Fig. 1), were excluded from further analysis. These traces could furthermore be identified by a markedly increased off-rate of H1 that results in a much lower affinity than expected from the experiments using freely diffusing molecules. The trajectories of immobilized nucleosomes binding to H1 free in solution (**Fig. 1e**) exhibited transitions between two states: the H1-bound nucleosome with high FRET efficiency and the unbound nucleosome with low FRET efficiency, in accordance with the free-diffusion experiments (**Fig. 1b,c**). We therefore analyzed the system in terms of a two-state model described by a rate matrix, ***K***, with transitions between unbound (N) and bound (HN) nucleosome:

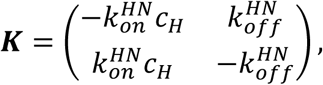

where *c*_H_ is the concentration of H1 in solution, and 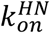 and 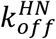 are the association and dissociation rate coefficients, respectively. The photon detection rates for donor and acceptor are estimated by applying maximum likelihood (MLH) analysis to the binned time traces^73^, where the likelihood, *L*_*m*_, for time trace *m* with bin size Δ, number of bins, *T*_*m*_, and (*N*_D,t_, *N*_A,t_) photons detected in time bin *t* is calculated according to

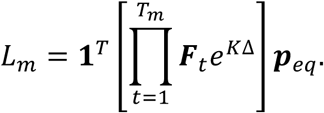

The population vector ***p***_*eq*_ describes the equilibrium distribution of states, for which ***Kp***_*eq*_ = 0 *1*^*T*^***p***_*eq*_ = 0. **1**^*T*^ = (1 1 …) is the transposed vector of ones. ***F***_*t*_ is a diagonal matrix with elements

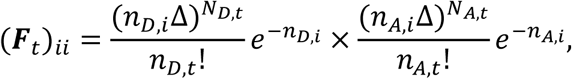

assuming Poisson statistics for the number of photons per bin. *n*_D,i_ and *n*_A,i_ are the mean photon detection rates of the *i*th state in the donor and acceptor channels, respectively. The photon detection rates and transition rate coefficients are then found by maximizing the sum of the logarithms of the likelihoods, Σ_*m*_ ln(*L*_*m*_) Using the Viterbi algorithm^74,75^, we next identify the most likely state trajectory using the photon detection rates and transition rate coefficients determined from the MLH analysis, and obtain the distributions of dwell times, which yielded rate coefficients in good agreement with those obtained with the MLH analysis when fit with single-exponential decays.

For describing the kinetics in the presence of ProTα, we assume that the nucleosome can additionally be in complex with both H1 and ProTα, forming the ternary complex PHN. Accordingly, we describe the measurements (Fig. 2b) with a three-state model (including N, HN, and PHN) based on the rate matrix

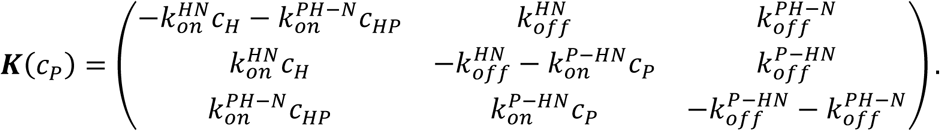

Even though a unique determination of all four additional rate coefficients besides the independently known 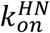 and 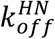 is not feasible based on the available data, the dependence of the observed transition rates between low- and high-FRET states on ProTα concentration can be described well with this model (solid lines in Fig. 2b), in contrast to a model without PHN (dashed lines in Fig. 2b), showing that the formation of the ternary complex PHN can explain the observed kinetic behavior. Interestingly, the simulations suggest that N and PHN exhibit similar inter-dye distances. It is noteworthy that the experimental data (Fig. 2b) can indeed only be described if we assume that HN exhibits high FRET efficiency and that both N and PHN exhibit low FRET efficiency, suggesting that ProTα binding leads to the opening of the linker DNA in the H1-nucleosome complex. Intermolecular FRET experiments (Supporting Information Fig. 3) demonstrate, however, that H1 dissociation from the nucleosome in the presence of ProTα occurs on a similar timescale as the transitions from high to low ⟨*E*⟩ (Fig. 2), indicating that nucleosome opening is coupled to rapid dissociation of H1 with ProTα.

### Simulations

#### Protein model

Each residue of ProTα and the core and linker histones was represented as a bead (C-beads) mapped on the Cα atom of the X-ray crystal structure (PDB ID: 5NL0). The potential energy had the following functional form^5^:

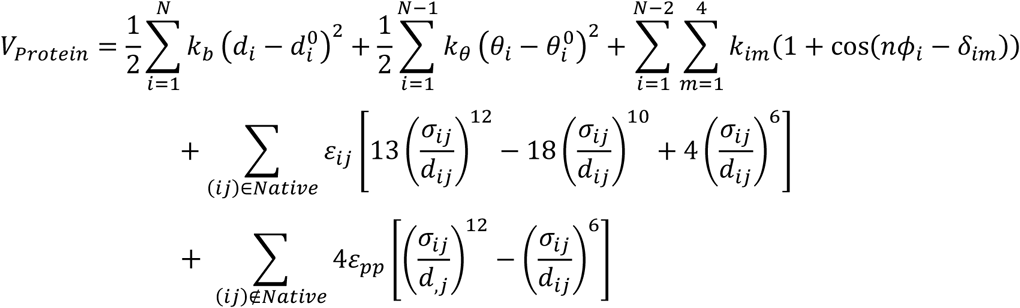

The first and second terms describe bonds and angles, respectively, using harmonic potentials with equilibrium values for bonds 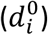 and angles 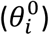 assigned according to the distances and angles between Cα atoms in the all-atom structure, as previously described^76^; values for unstructured regions were taken from an extended conformation The third term defines the torsions (dihedrals) between four beads linked by three bonds. The dihedral parameters were obtained via a knowledge-based potential based on the protein data bank^76^. The fourth term describes electrostatic interactions between charged residues using a screened Coulomb potential. Aspartate and glutamate residues were assigned a charge of −1, lysine and arginine a charge of +1, and histidine a charge of +0.5 to account for the pK_a_ of histidine of ∼6.5. A charge of 0 was assigned to all the other beads. In the Coulomb term, *q* is the charge of residue *i*; *ε*_*O*_ is the permittivity of free space; and *ε*_*d*_ is the relative dielectric constant of water, which was set to a value of 80. The Debye screening length, *λ*_*D*_, is given by

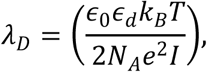

where *N*_*A*_ is Avogadro’s constant, *e* is the elementary charge, and */* denotes the ionic strength, *k*_*B*_ is the Boltzmann constant and *T* is temperature.

The fifth and sixth terms define short-range attractive interactions between beads. These interactions are applied differently to disordered and folded regions of the proteins. For the folded regions, native interactions are enforced using a 12-10-6 pair potential^76^. The values of *ε*_*i,j*_ were calculated based on the globular domain of H1 and core histones according to the native-centric model of Karanicolas and Brooks^76^. *σ*_*ij*_ was set according to the Cα distances in the crystal structure. For disordered regions and interactions between non-native residue pairs and the folded regions (i.e., those not in contact in the native state), the *ε*_*pp*_ term defining the strength of the interaction between beads was tuned to the value of 0.16 *k*_B_*T* (∼0.40 kJ mol^-1^), the value which was shown to produce the best agreement with the experimentally determined FRET efficiencies (Fig. 3) for the 1:1 ProTα/H1 complex^5^, while the value of *σ*_*ij*_ *= (σ*_*i*_ *+ σ*_*j*_)/2 was determined based on *σ*_*i*_ specific to each residue type^77^.

#### DNA model

The coarse-grained model of the DNA featured three beads representing the phosphate (P), ribose (R), and base (B) moieties of a nucleotide and were mapped to the P, C4’, and N1 atoms from the full atom DNA structure, respectively. Phosphate beads were assigned a charge of −1. The functional form of the potential for the DNA was:

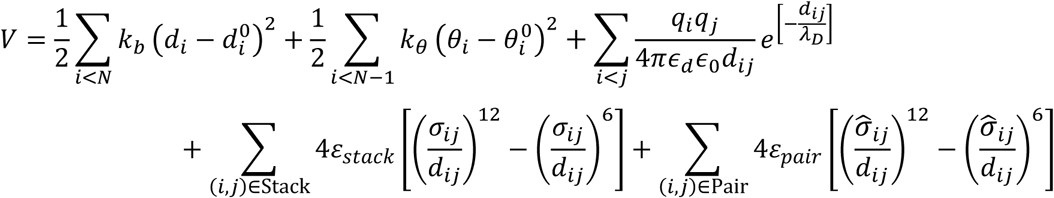

The first two terms describe bonds and angles between beads, with parameters chosen to reproduce the equilibrium structure of B-DNA. A screened Coulomb potential was used for electrostatic interactions as described above. Non-bonded Lennard-Jones potentials were used to reproduce pairing and stacking between bases. The values of the *ε*_*stack*_ and *ε*_*pair*_ were set to 3.0 *k*_B_*T* (∼7.5 kJ mol^-1^) and 3.5 *k*_B_*T* (∼8.8 kJ mol^-1^), consistent with previously estimated free energy values for base stacking^78^. This is the same model used previously to describe a DNA hairpin^50^; only the base-pairing energy, *ε*_*pair*_, was optimized to obtain the persistence length (∼50 nm) characteristic of double-helical DNA^78^.

#### Protein-DNA interaction potential

The potential energy for the interactions between DNA and protein beads is composed of a screened Coulomb potential, which describes the interaction between charged beads, and short-range potentials:

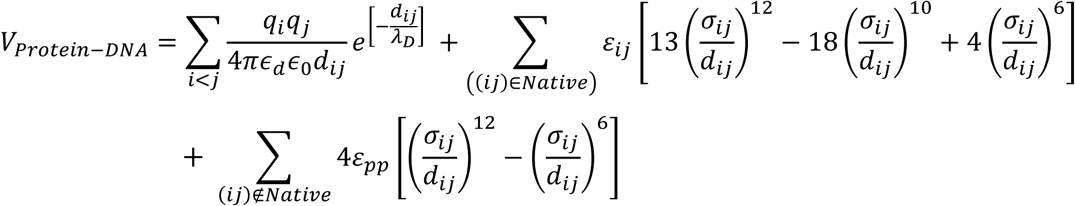

In this case, pairs of beads interacting via native interactions were identified based on the crystal structure of the nucleosome (pdb code: 5NL0). Native interactions were defined between protein beads and ribose (nucleobase) beads if any heavy atom of an amino acid falls within 0.5 nm from any heavy atom of that ribose (nucleobase). *ε*_*ij*_ for these interactions was set to 2 *k*_B_*T* (∼5 kJ mol^-1^) to ensure these interactions remained formed. For the interactions between the H1 globular domain and the DNA dyad^79,80^, we reduced the native *ε*_*ij*_ to allow for globular domain fluctuations and identified the interaction strength that yielded the best agreement between experimental and computed FRET efficiencies, yielding a value of 1 *k*_B_*T* (∼2.5 kJ mol^-1^). For non-native interactions between the disordered regions of the proteins (histone tails and ProTα) and the DNA, *ε*_*pp*_ was set to 0.06 *k*_B_*T* (∼0.15 kJ mol^-1^) and *a*_*ij*_ to 0.6 nm. The values of *ε*_*ij*_ and *ε*_*pp*_ were chosen to optimize the agreement between simulated and experimentally obtained FRET efficiencies. Note that the optimal strength of non-native interactions, *ε*_*pp*_, is the same as that obtained for a non-specific complex between a protein chaperone and DNA hairpin^50^; only the native contact parameters representing specific interactions needed to be tuned for this system. For simulations with the H1 globular domain rotated by 180° on the nucleosomal dyad, a coordinate transformation was performed and steric clashes relaxed by energy minimization. Native interactions between the rotated globular domain and the DNA were retrieved from the rotationally symmetric list of contacts between the globular domain and the DNA in its native conformation. A summary of all parameter values defining bonded and non-bonded interactions in the models is provided in Supporting Information Tables 3 and 4. In summary, the similarity of the parameters to those used in previous work is testament to their transferability. The parameters that were tuned in this work were (i) the DNA base-pairing energy *ε*_*pp*_ and (ii) the strength of the specific, native protein-DNA interactions.

#### Replica-exchange Langevin dynamics simulations of the nucleosome

Langevin dynamics simulations were performed using a modified version of GROMACS 5.4.1^81,82^. The nucleosome was placed at the center of a box of 30 nm x 40 nm x 30 nm and simulated with and without H1 bound using periodic boundary conditions and charge screening equivalent to 165 mM salt. After energy minimization, 33 replicas were simulated in a temperature range between 298.15 and 648.23 K. Exchange attempts between conformers of neighboring replicas in temperature space were made every 5 ps and accepted according to the Metropolis criterion. Each replica was run for 1 μs, using a timestep of 10 fs and a friction coefficient set at 0.2 ps^-1^. Particle velocities for each replica were randomly seeded according to a Boltzmann distribution. Interaction energies were calculated in direct space using a cutoff of 2.5 nm, with neighbor-searching performed every 0.1 ps.

To test the influence of the FRET dyes on the simulation results, we performed a set of simulations in which the dyes were represented explicitly. Each dye and linker were described by five beads, reflecting the approximate equivalence to a five-amino acid chain^83^, linked by bonds with a length of 0.38 nm and bond angles of 110° (the dihedral term was omitted). The interaction between the dyes and the protein or DNA was characterized by a weakly attractive short-range interaction energy, with ε = 0.001 kJ mol^-1^ and σ = 0.6 nm. The fourth and the fifth beads carried a charge of −1 each to reflect the total charge of −2 of the Alexa fluorophores used in the measurements.

Mean FRET efficiencies (⟨*E*⟩) were calculated from simulations according to the interbead distance distributions, *P*(*r*), of the fluorescently labeled residues:

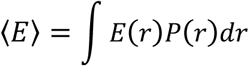

and the Förster equation:

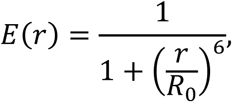

using the Förster radius, *R*_0_, of 5.4 nm of the Alexa Fluor 488/594 dye pair. The agreement between *⟨E⟩* from experiments and simulations was quantified by the concordance correlation coefficient^84^, *ρ*_*c*_, between the two datasets:

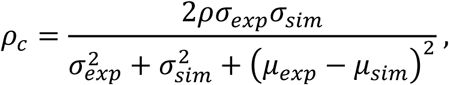

where *σ*_exp/sim_ indicate the standard deviations, *μ*_exp/sim_ the mean values, and *ρ* the linear (Pearson) correlation coefficient between the values from experiments and simulations. The concordance correlation coefficient is a stricter measure than the linear correlation coefficient: *ρ*_*c*_ is in general smaller than *ρ* and is equal to it only if *σ*_*exp*_ *= σ*_*sim*_ and *μ*_*exp*_ *= μ*_*sim*_.

#### Langevin dynamics simulations of the nucleosome in the presence of ProTα

Langevin dynamics simulations of the nucleosome were performed at a concentration of ProTα of 100 μM, i.e., within the experimental concentration range, by placing 10 molecules of ProTα and the nucleosome bound to H1 in a cubic box of 55 nm × 55 nm × 55 nm. To reduce the equilibration and the dissociation for the simulations with an external pulling force (see below) to an accessible time scale, we decreased the value of *ε*_pp_ between ProTα and core histone beads to ∼0.0025 kJ mol^-1^, while leaving *ε*_pp_ between ProTα and H1 beads at 0.6 kJ mol^-1^. Simulations of the interaction of ProTα with the H1-nucleosome complex were run at 298.15 K with the same H1-nucleosome parameters optimized to match the equilibrium FRET data. Different conformations of the ProTα-H1-nucleosome complex were isolated from these runs and used as starting points for the pulling simulations in the presence of ProTα, with identical simulations performed in the absence of ProTα.

Constant-force pulling simulations were performed at 11 forces from 30 to 130 pN, applied between the center of mass of the H1 globular domain and the center of mass of the beads of the DNA involved in native interactions with H1. At each force value, 30 replicates were performed. Simulations were stopped when full dissociation of H1 from the nucleosome was achieved (distance between any bead pairs of H1 and DNA >2.5 nm) or 10 μs was reached. We quantified the average escape times, *τ*, of H1 or the H1-ProTα complex from the nucleosome at each force from the escape times of individual runs, *τ*_*i*_, according to the maximum-likelihood estimator^85^:

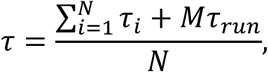

where *N* is the number of replicates where dissociation occurs within the simulation time, and *M* is the number of replicates where dissociation does not occur during the total allocated time per run of *τ*_*run*_ = 10 μs. To extrapolate to the escape times in the absence of force, we fitted *τ* as a function of the applied force, *F*, using the Dudko-Hummer-Szabo (DHS) model^86^:

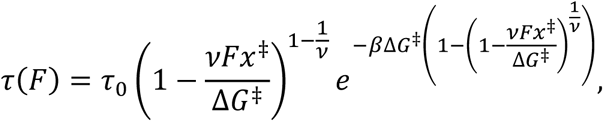

where *τ*_0_ is the escape time in the absence of force, *x*^‡^ the distance to the transition state, Δ*G*^‡^ the apparent free energy of activation in the absence of an external force, and *v* is an exponent specifying the shape of the free-energy profile, with *v* = 1/2 for a harmonic well with a cusp-like barrier, *v* = 2/3 for a potential with both linear and cubic terms, and *v* = 1 for Bell’s model^87^. The data were fitted using all three exponents, with *τ*_0_, Δ*G*^‡^, and *x*^‡^ as fit parameters. Errors of the fit parameters were estimated by bootstrapping: 1000 data sets were generated by drawing from a Gamma distribution of the *τ* value, with the shape parameter of the Gamma distribution set to the number of runs where full dissociation of H1 from the nucleosome occurred, and the scale parameter set to the mean dissociation time of H1. We then fitted each data set using the DHS model. Errors are the standard deviations of the resulting distributions of fit parameter values reported in Supporting Information Table 4.

## Supporting Information

**Supporting Information Figure 1.**
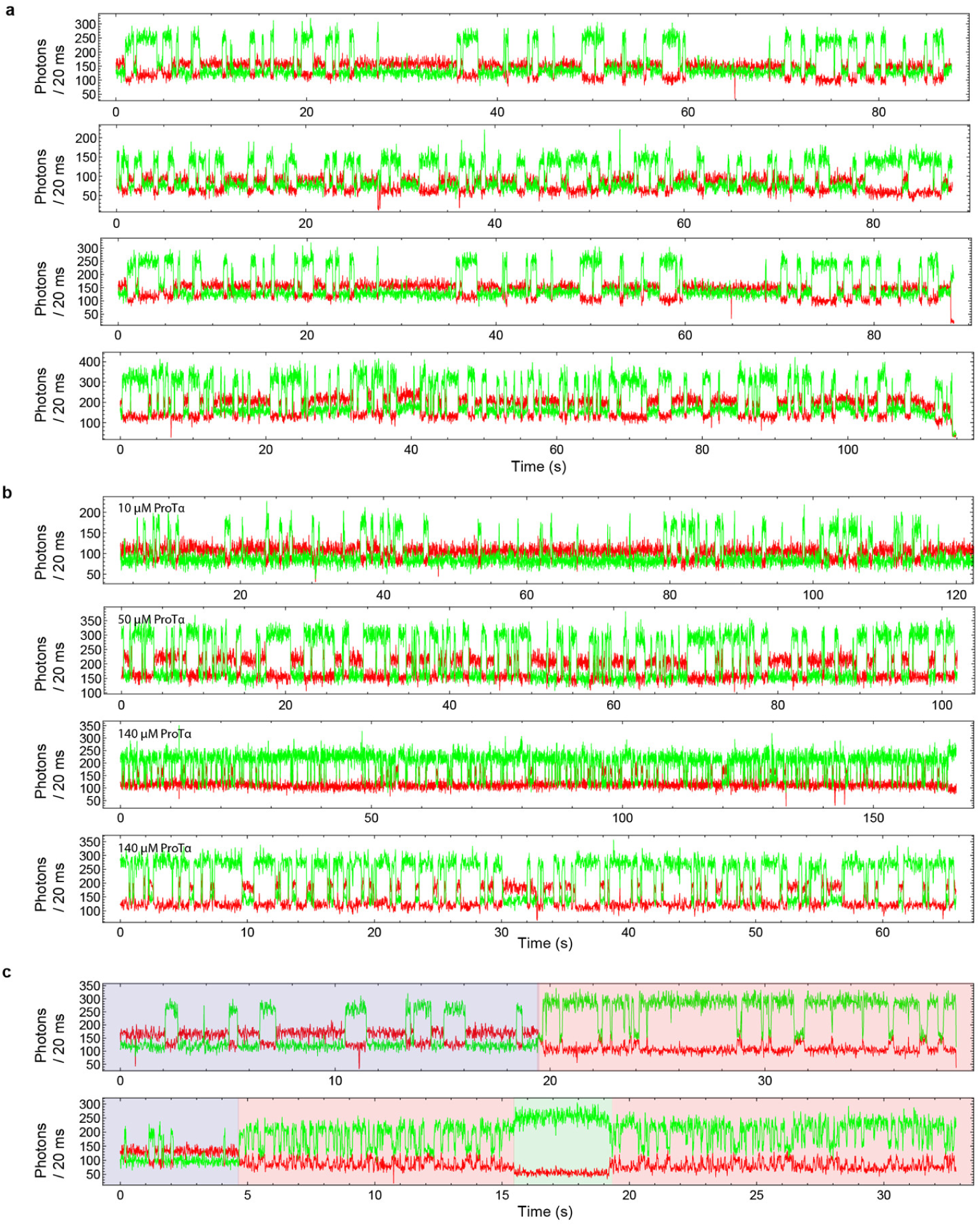
Fluorescence time traces of surface-immobilized nucleosomes. **a**, Representative time traces of unlabeled H1 binding to double-labeled nucleosomes, at 360 mM ionic strength and 2 nM H1 in the absence of ProTα. **b**, Representative time traces of H1 binding to nucleosomes in the presence of 10, 50, or 140 µM ProTα, at 340 mM ionic strength and 3 nM H1. **c**, Representative time traces showing unwrapping of nucleosomes. Unwrapping occurs after ∼20 s in the upper trace (red-shaded range). Such partially wrapped states, which have previously been observed, may consist of one or both linker DNA arms partially detached from the nucleosome core^27,28^; corresponding segments or the entire traces were excluded from the analysis (see Methods). Binding events to these partially wrapped states are still observed, consistent with reports that H1 retains high affinity to nucleosomes that contain only one linker DNA arm^23^. The lower trace shows a further unwrapping event (15.5-19 s, green-shaded range) that displays no anti-correlated changes in FRET before returning to a binding-competent partially unwrapped state.

**Supporting Information Figure 2.**
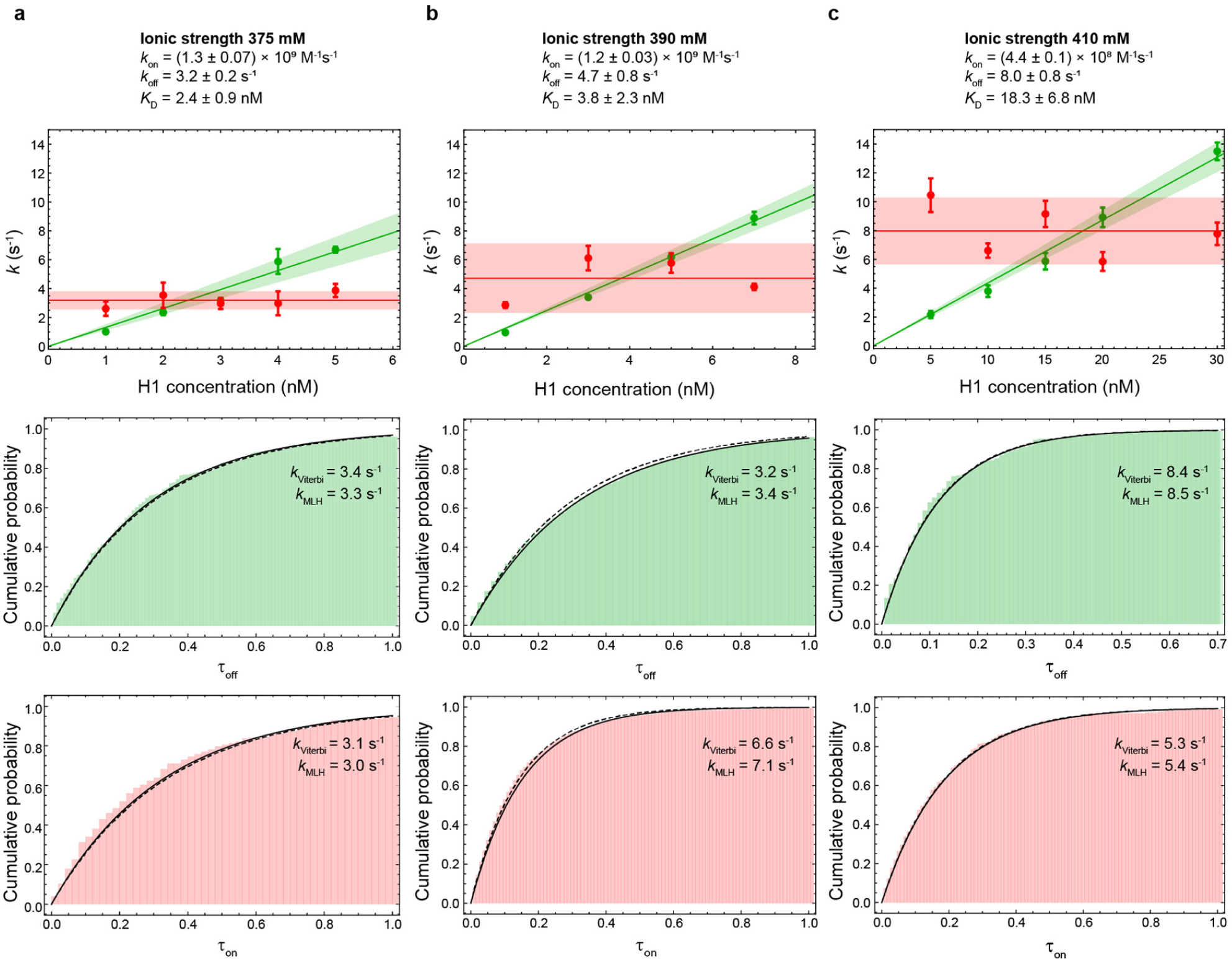
H1-nucleosome binding kinetics at different ionic strengths. Association (green) and dissociation (red) rates of the H1-nucleosome complex as a function of H1 concentration, at (**a**) 375 mM, (**b**) 390 mM, and (**c**) 410 mM ionic strength, and corresponding examples of cumulative dwell-time probability distributions, at 2 nM, 3nM, and 20 nM H1, respectively. The rate coefficients and equilibrium dissociation constants resulting from the fits (lines) are shown in each panel. Error bars show one standard deviation estimated from ten bootstrapping trials; shaded areas indicate 95% confidence intervals. The dwell times determined with the Viterbi algorithm are displayed as cumulative probability distributions and fits with a single-exponential function to yield the transition rates (solid lines). A single-exponential function based on the transition rates determined by MLH analysis is plotted for comparison (dashed lines) to illustrate the agreement with the rates determined by Viterbi analysis.

**Supporting Information Figure 3.**
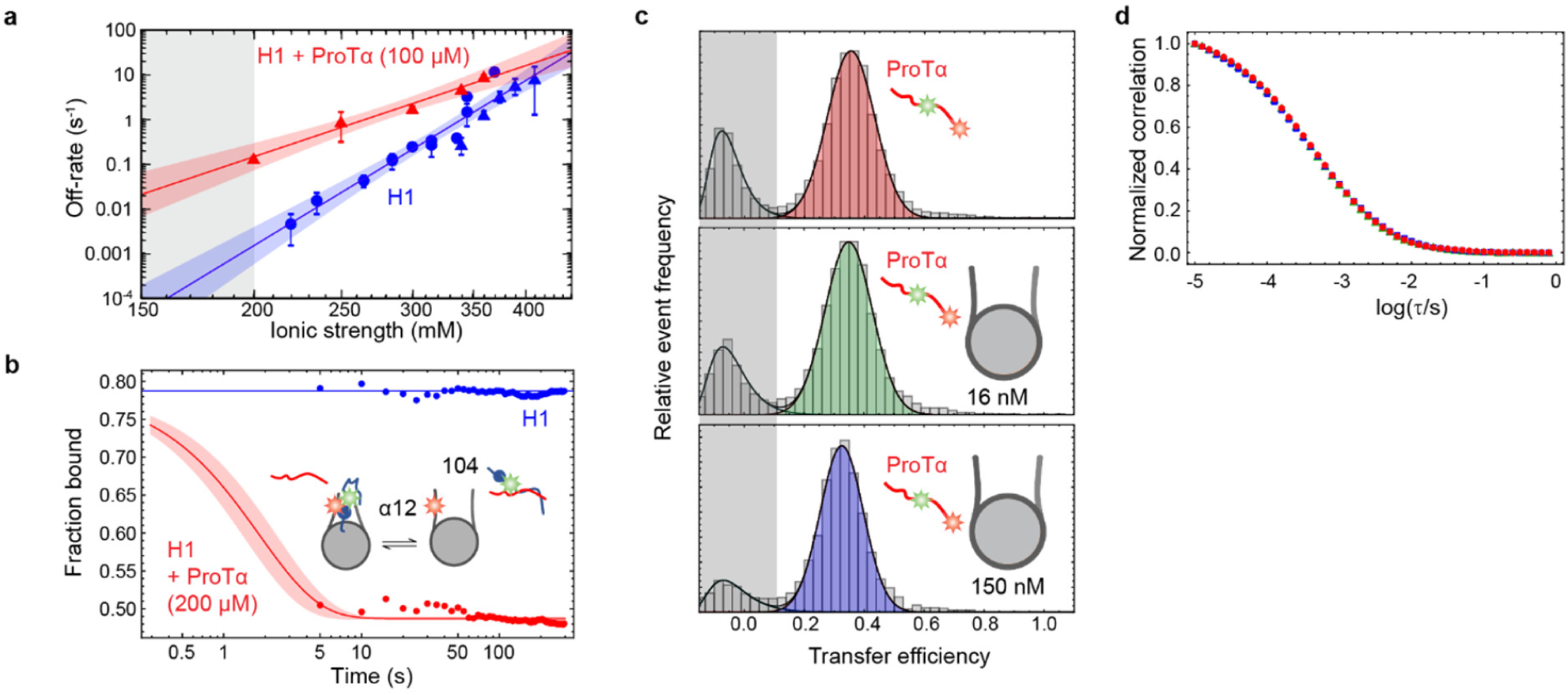
ProTα accelerates H1 dissociation from the nucleosome. Effects of ProTα on the H1-nucleosome complex. **a**, Observed off-rates of H1 from nucleosomes in the absence of ProTα (blue triangles: surface experiments, blue circles: from *K*_D_ and diffusion-limited on-rate coefficient) and transition rate from high to low FRET efficiency (*E*_high_ → *E*_low_, red triangles) in the presence of 100 µM ProTα, as a function of ionic strength with fits to the Lohman-Record model^59^ for extrapolation into the physiological ionic strength range (gray shading). **b**, Fraction of donor-labeled H1 (labeled with Alexa Fluor 488 in position 104) bound to acceptor-labeled nucleosome (labeled with Alexa Fluor 594 at position α12), based on transfer efficiency histograms from time-resolved intermolecular FRET experiments on freely-diffusing molecules, in the absence (blue) and presence (red) of 200 µM ProTα, at 200 mM ionic strength. A transfer efficiency histogram was constructed every 5 s after mixing of ProTα (dead time 5 s). A fit of the data shows that ProTα accelerates dissociation of H1 from the nucleosome (fraction bound is reduced from ∼0.8 to ∼0.5), resulting in a dissociation rate of at least 0.6 ± 0.1 s^-1^, much faster than the value of ∼10^−3^ s^-1^ expected in the absence of ProTα (panel **a**). **c**, Transfer efficiency histograms of fluorescently-labeled ProTα (with Alexa Fluor 488 and Alexa Fluor 594 in positions 56 and 110), either in isolation (top) or in the presence of 16 nM (middle) and 150 nM (bottom) unlabeled nucleosomes, and at 165 mM ionic strength. Negligible differences in average transfer efficiency are observed between the histograms, and for the bottom histogram can be explained by considerably higher background fluorescence which decreases the apparent transfer efficiency slightly. For comparison, when bound to H1, the transfer efficiency of this ProTα variant increases from 0.36 to 0.54^5^. **d**, Donor-acceptor cross-correlation curves were used to compare the translational diffusion of ProTα in isolation and in the presence of nucleosomes, at 165 mM ionic strength (same color scheme as in panel b). Binding of ProTα to the much larger nucleosome particles would be expected to substantially increase its diffusion time. The measured diffusion time of ProTα in the presence of nucleosomes is, however, almost identical to its diffusion time in isolation. The data in panels c and d indicate that, without H1 present, a direct interaction between ProTα and nucleosomes does not take place at these protein concentrations.

**Supporting Information Figure 4.**
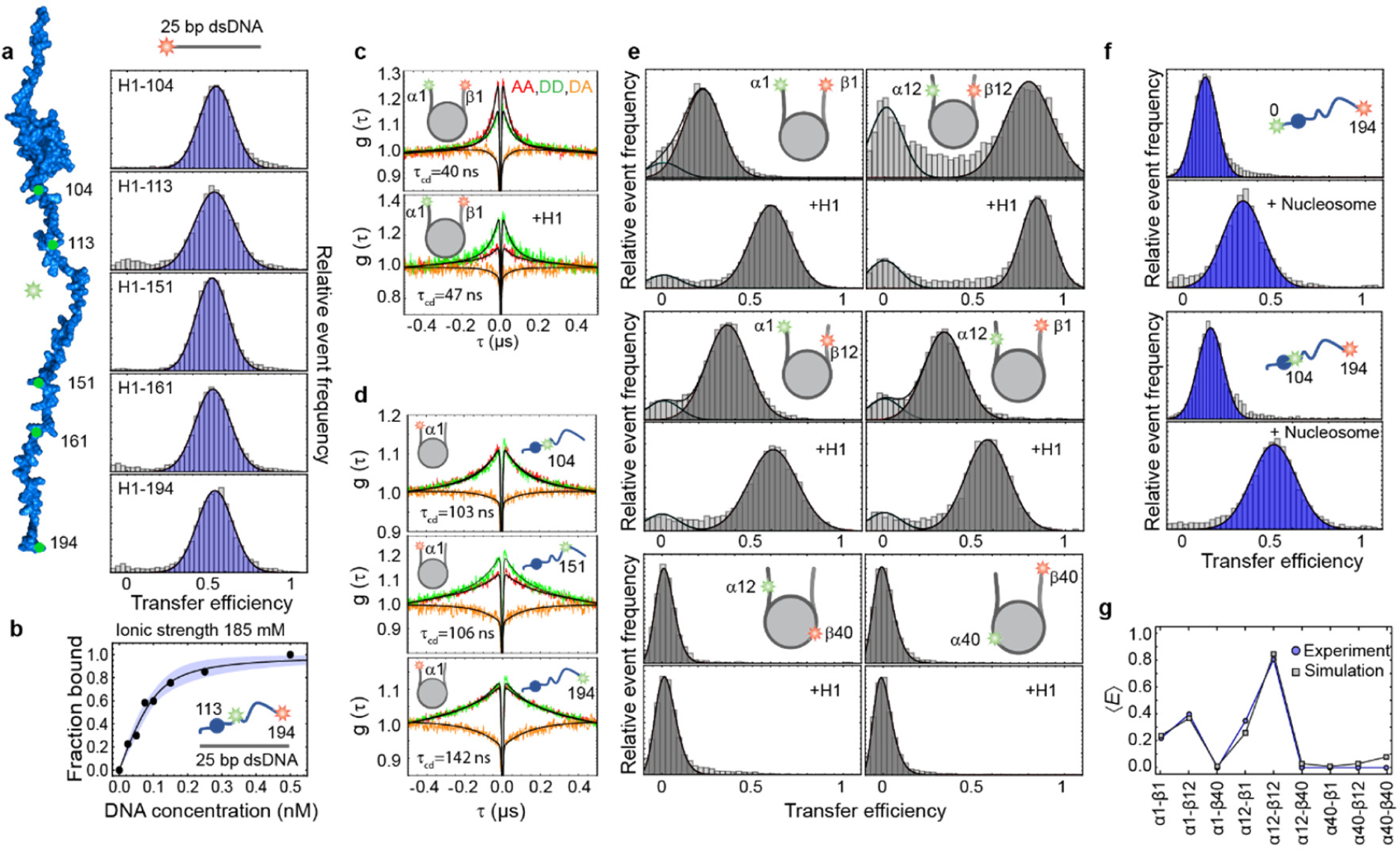
Interactions and dynamics of H1, nucleosomes, and DNA. **a**, Transfer efficiency histograms from intermolecular FRET experiments between donor-labeled H1 and acceptor-labeled 25-bp dsDNA with a sequence corresponding to the α-linker used in the nucleosome constructs (see Supporting Information Table 2). **b**, Binding isotherm for binding of double-labeled H1 to 25-bp dsDNA at 185 mM ionic strength. The affinity of H1 for DNA (∼25 ± 2 pM) is two orders of magnitude lower than for the nucleosome at the same ionic strength (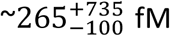, Fig. 1d). **c**, Nanosecond fluorescence correlation spectroscopy (nsFCS) of nucleosomes labeled at the α1 and β1 positions, without (upper) and with (lower) unlabeled H1 bound; both exhibit fast dynamics of the linker DNA arms. **d**, Intermolecular nsFCS of H1 donor-labeled in positions 104, 151, and 194 and nucleosomes acceptor-labeled at position α1 shows that rapid dynamics of H1 relative to the linker DNA are present throughout the C-terminal region of H1 in complex with the nucleosome. **e**, Intramolecular transfer efficiency histograms of nucleosomes with FRET dyes at different positions on the α and β strands, in the presence and absence of saturating concentrations of unlabeled H1. Changes in transfer efficiency report on changes in linker DNA arrangement upon H1 binding. For variants α1/β1, α1/β12, α12/β12, and α12/β1, the peaks at *E* ≈ 0 (light gray) are incompatible with a correctly formed nucleosome and were thus excluded from the analysis. **f**, Intramolecular single-molecule FRET experiments with dyes at different positions along the H1 sequence and unlabeled nucleosomes. Labeling positions are indicated as numbers and arrows along the structural or cartoon representations of H1. Experiments were conducted at 165 mM ionic strength unless stated otherwise. **g**, Comparison between average intra-nucleosome FRET efficiencies from experiment (circles) and simulation (squares), in the absence of H1. The excellent agreement attests to the quality of the simulation model for describing the nucleosome structure.

**Supporting Information Figure 5.**
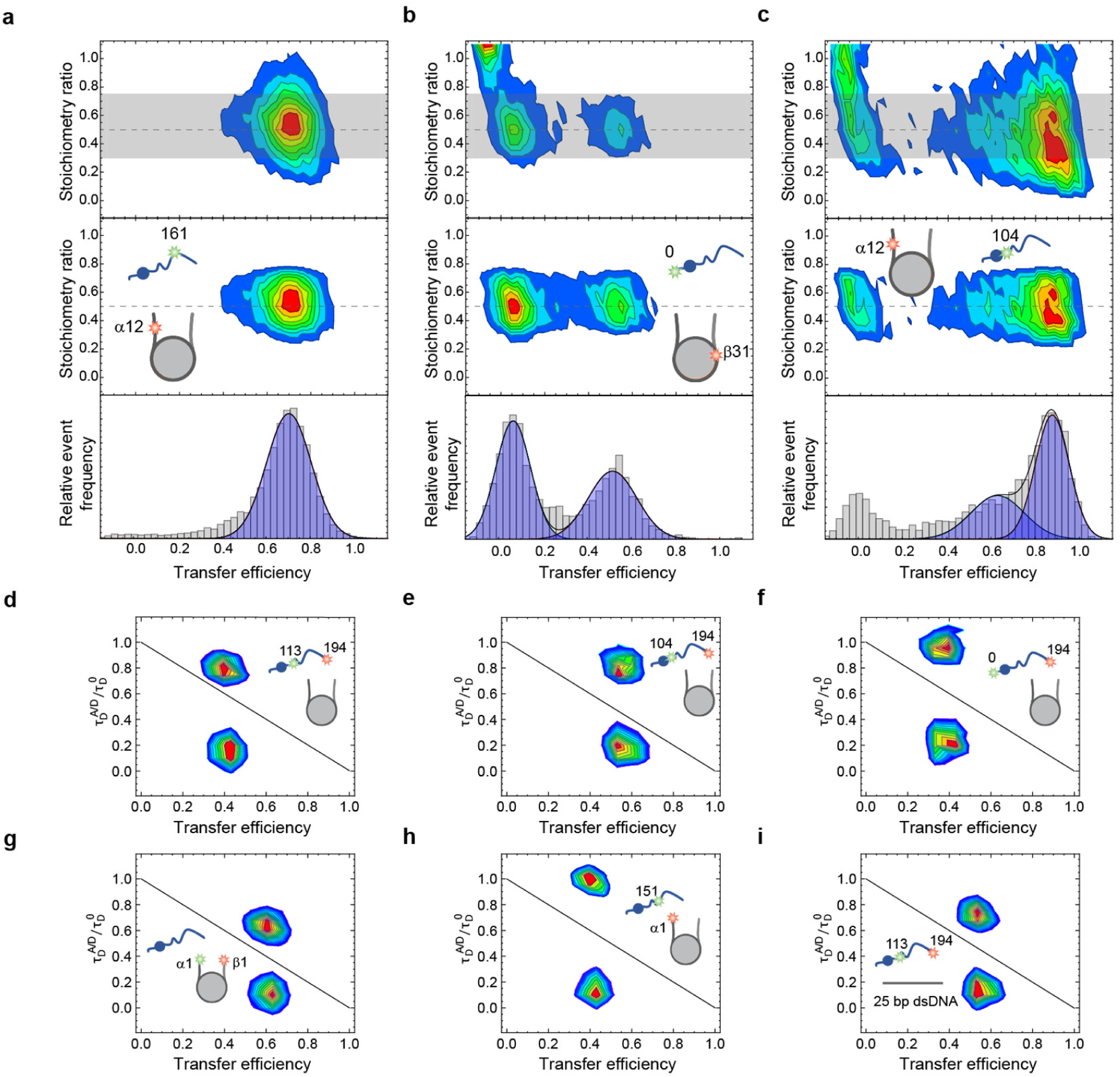
Representative examples of stoichiometry measurements and fluorescence lifetime analysis of H1-nucleosome and H1-DNA complexes. **a-c**, 2D-histograms of stoichiometry ratio vs. transfer efficiency from intermolecular FRET between donor-labeled H1 and acceptor-labeled nucleosome using pulsed interleaved excitation. The labeling positions are indicated in the schematics above the graphs. The gray-shaded areas indicate the stoichiometry ranges used for filtering out donor-only fluorescence bursts. A 1:1 complex results in a stoichiometry ratio of 0.5 (dashed horizontal line). Most of the transfer efficiency histograms in Fig. 3 contain a single population, as in (**a**). In some cases, such as in (**b**), a clear second population is observed that is well separated from the donor-only population. In other cases, i.e. in (**c**), an additional residual population at a transfer efficiency close to zero can remain even after filtering due to a large signal from molecules lacking an active acceptor dye. **d-i**, Fluorescence lifetime analysis. Plots showing the fluorescence lifetimes of Alexa Fluor 488 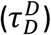 and Alexa Fluor 594 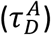 normalized by the intrinsic donor lifetime 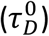 as a function of transfer efficiency. If fluctuations in transfer efficiency occur on a timescale between the donor fluorescence lifetime and the burst duration (∼4 ns-1 ms), the normalized donor and acceptor lifetimes cluster above and below the static FRET line (solid diagonal line)^70,71^, respectively. The substantial deviations from the diagonal indicate a broad distribution of distances in all samples, as previously observed for intrinsically disordered proteins^47,88^. **d**, H1^113-194^ + unlabeled nucleosome; **e**, H1^104-194^ + unlabeled nucleosome; **f**, H1^0-194^ + unlabeled nucleosome; **g**, Nucleosome^α1-β1^ + unlabeled H1; **h**, H1^151^ + nucleosome^α1^; **i**, H1^113-194^ + 25 bp dsDNA.

**Supporting Information Figure 6.**
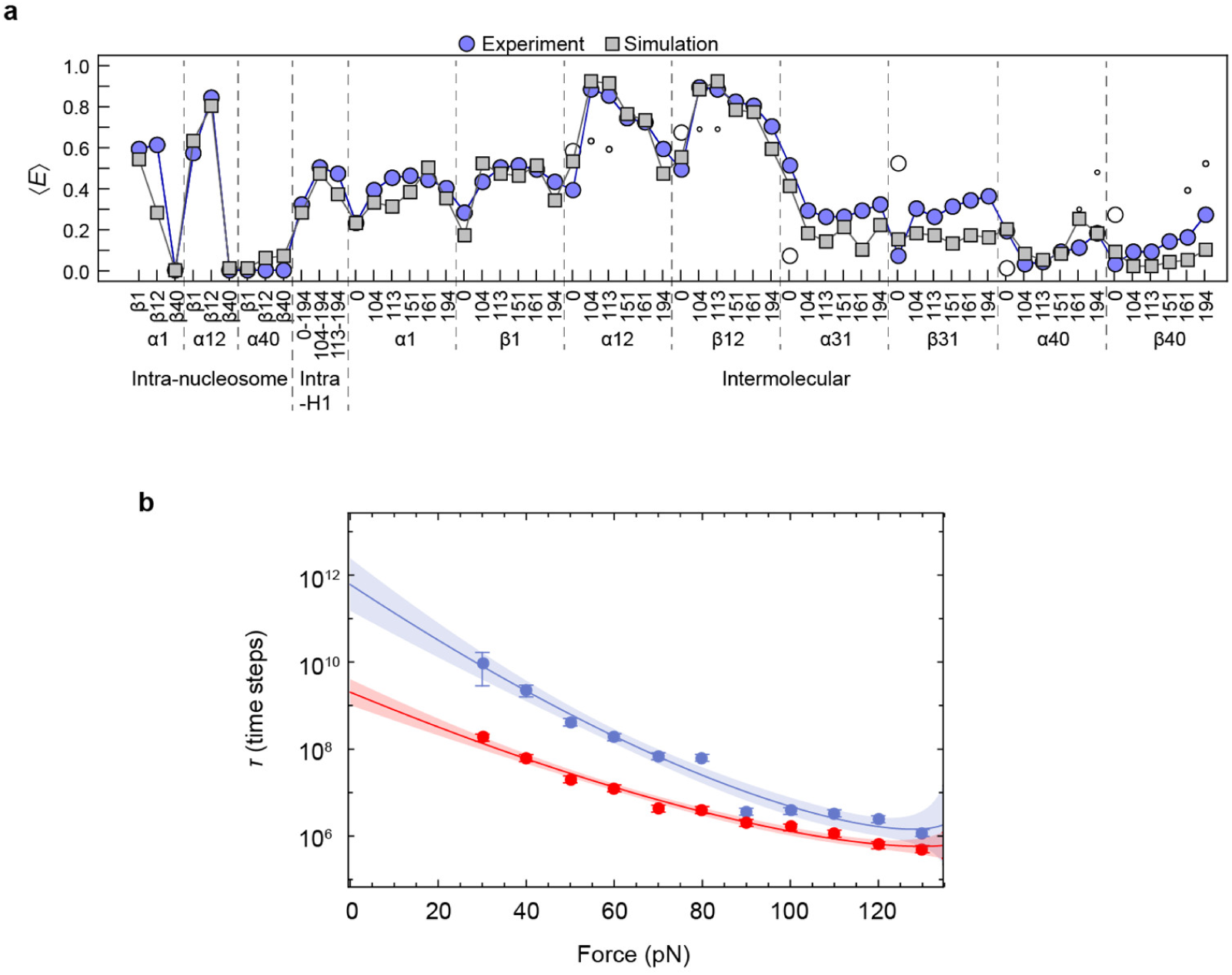
Additional simulation data. **a**, Comparison between FRET efficiency values from experiment and simulation using an explicit representation of dyes. Experimental FRET efficiency values (purple and empty circles) are identical to the ones shown in Fig. 3. FRET efficiency values from simulation (gray squares) are computed from simulations where a representation of the dyes was included in the corresponding labeling positions (see Methods). **b**, Alternative analysis of the mean escape times (*τ*) of H1 from the nucleosome as a function of the pulling force. The results in the absence and presence of ProTα are shown in blue and red, respectively. The solid lines show the fit of the mean escape times using the Dudko-Hummer-Szabo model^89^ using an exponent of 2/3 (rather than 1/2, Fig. 4). Even though the resulting fit is slightly worse than with an exponent of 1/2 (Fig. 4, Supporting Information Table 4), the acceleration of dissociation by ProTα extrapolated to zero force (*τ*_0_) is similar, illustrating the robustness of the analysis. Note that the Bell model (v = 1) is not suitable for describing the data as it fails to describe the decreasing slope at higher force, associated with a shift of the barrier on the energy landscape.

**Supporting Information Table 1.** Binding parameters of H1-nucleosome interactions. Equilibrium and rate constants for nucleosome-H1 interactions at different ionic strengths, in the absence and presence of 100 µM ProTα. The equilibrium dissociation constants are either from the ratio of the rate coefficients based on the surface experiments 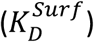 or from the binding isotherms measured on freely diffusing molecules 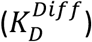. The agreement between the two methods confirms that surface immobilization does not interfere with the interactions between nucleosomes, H1, and ProTα. Rate constants were determined via maximum-likelihood analysis of fluorescence time traces collected with surface-immobilized nucleosomes in the presence of 0.5-30 nM of unlabeled H1. Errors on rate constants and 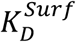 are based on ten bootstrapping trials errors from global likelihood maximization. For experiments on freely diffusing H1-nucleosome complex with 100 µM ProTα, H1 was titrated into a solution containing nucleosome until binding was saturated. Errors are based on pipetting error estimates.

| H1-nucleosome complex | | | | | H1-nucleosome complex + 100 μM ProTα | | |
| --- | --- | --- | --- | --- | --- | --- | --- |
| ionic strength<br>(mM) | $K_D^{Diff}$<br>(nM) | $K_D^{Surf}$<br>(nM) | $k_{on}$<br>($M^{-1}s^{-1}$) | $k_{off}$<br>($s^{-1}$) | ionic strength<br>(mM) | $K_D^{Diff}$<br>(nM) | $K_D^{Surf}$<br>(nM) |
| 360 | $1.7 \pm 0.1$ | $0.9 \pm 0.3$ | $(1.5 \pm 0.04) \cdot 10^9$ | $1.3 \pm 0.1$ | 200 | $0.05 \pm 0.02$ | $0.35 \pm 0.03$ |
| 375 | $3 \pm 0.2$ | $2.4 \pm 0.9$ | $(1.3 \pm 0.07) \cdot 10^9$ | $3.2 \pm 0.2$ | 250 | $0.16 \pm 0.02$ | $0.76 \pm 0.05$ |
| 390 | $5 \pm 0.3$ | $3.8 \pm 2.3$ | $(1.2 \pm 0.03) \cdot 10^9$ | $4.7 \pm 0.8$ | 300 | $0.95 \pm 0.2$ | $2.45 \pm 0.07$ |
| 410 | $9.7 \pm 0.6$ | $18.3 \pm 6.8$ | $(4.4 \pm 0.01) \cdot 10^8$ | $8.0 \pm 0.8$ | 340 | $3.15 \pm 0.5$ | $8.48 \pm 0.06$ |
| | | | | | 360 | $10.7 \pm 0.9$ | $24.4 \pm 1.1$ |

**Supporting Information Table 2.**
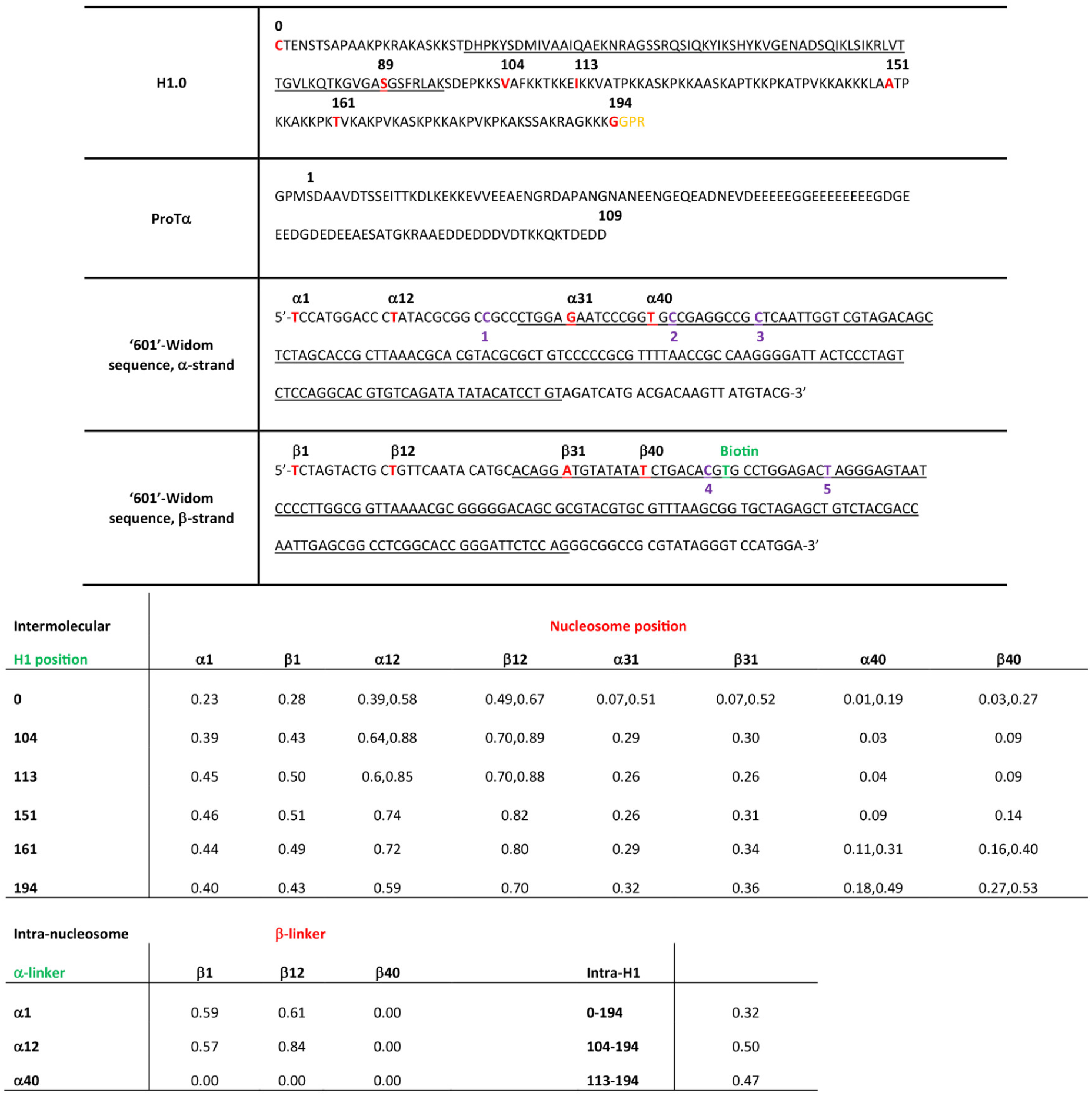
Sequences of H1, ProTα, and nucleosome DNA, and transfer efficiencies of all labeling pairs. Upper table: Sequences of H1 and ProTα constructs and the 601-Widom sequence^21^ used for nucleosome reconstitution. Residues and nucleotides in red indicate fluorophore labeling sites. H1 residues in yellow remained after proteolytic cleavage of the hexahistidine tag with thrombin. The underlined segment of H1 represents the globular domain^5^. The nucleotide indicated in green was modified with a biotin linker for surface immobilization of the nucleosomes. Underlined nucleotides constitute the 147-bp 601-Widom sequence, which forms the core nucleosome particle; the remaining nucleotides correspond to the linker DNA^90^. For DNA labeling, thymine modified with a C6-aminolinker was incorporated for the reaction with the succinimidyl ester of the fluorescent dye. Purple labels (1-5) denote the 3’- end of oligonucleotide primer used to PCR-amplify the corresponding nucleosomal DNA, either with or without previous fluorescent labeling (1: unlabeled and α1−labeled nucleosomes, 2: α12/α31-labeled nucleosomes, 3: α40-labeled nucleosome, 4: β1/β12/β31/β40-labeled nucleosomes, 5: biotinylated nucleosomes). In all cases, the 5’-end nucleotide of the respective PCR primer is the one shown at the 5’-end of the Widom sequence. Lower table: Transfer efficiencies of all labeling pairs measured in this study. For intermolecular and intra-nucleosome FRET efficiencies, the donor and acceptor dye positions are indicated in green and red, respectively. In cases where two peaks were observed, both values are given. The accuracy of the transfer efficiency values is approximately ±0.04^72^, the precision approximately ±0.02.

**Supporting Information Table 3.**
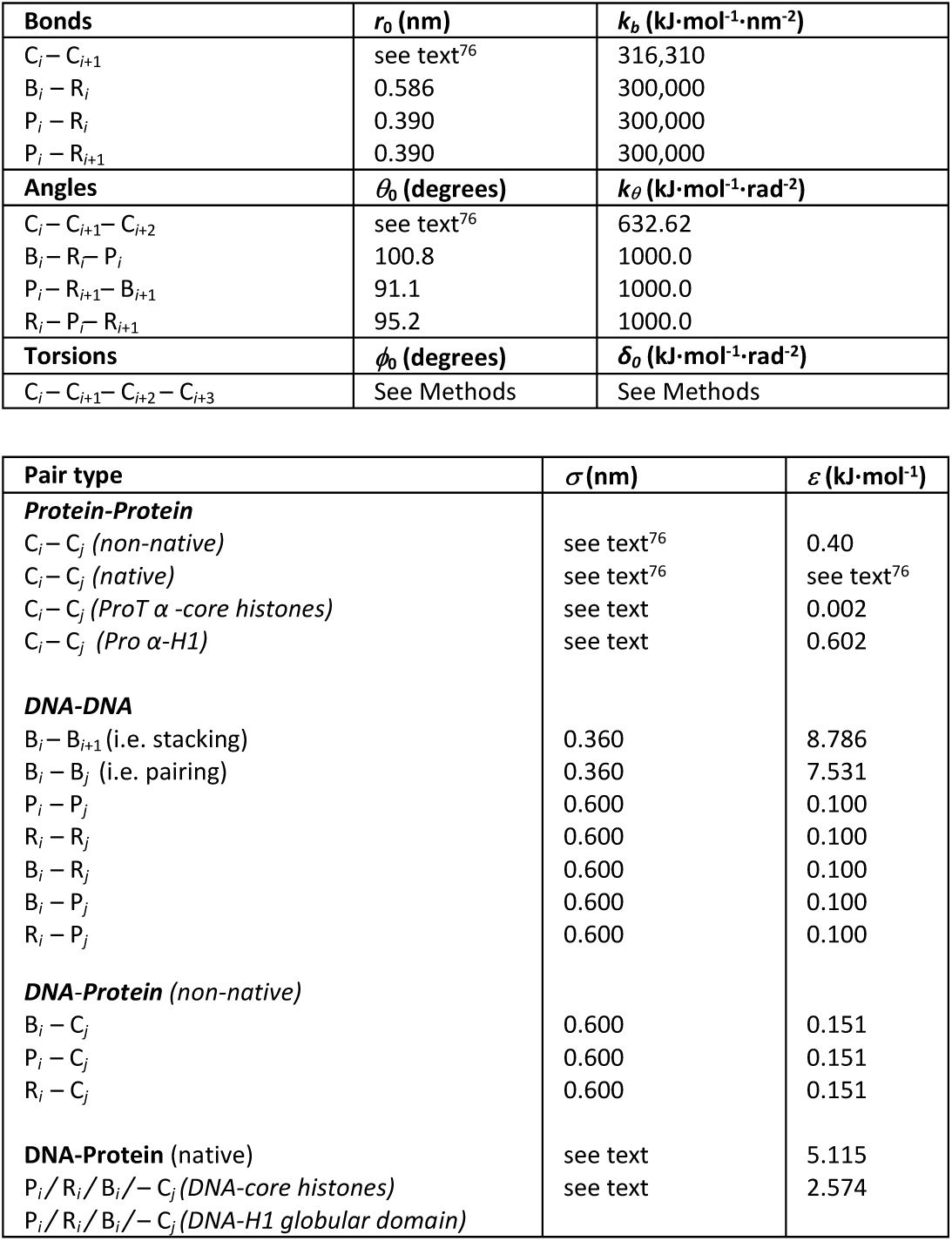
Bonded and non-bonded interactions between protein and DNA beads used for simulations. List of bonded and non-bonded parameters describing the interaction between protein and DNA beads. Details for some parameters are given in the text.

**Supporting Information Table 4.** Results of the fit of pulling simulations.

|  | *Scaling factor ($\nu$)* |  |
| --- | --- | --- |
|  | *1/2* | *2/3* |
| *Simulations in the absence of ProTα* |  |  |
| $\tau_0$ (time steps) | $(13 \pm 0.9) \cdot 10^{11}$ | $(6.1 \pm 0.9) \cdot 10^{11}$ |
| $x^\ddagger$ (nm) | $0.78 \pm 0.08$ | $0.65 \pm 0.08$ |
| $\Delta G^\ddagger$ ($k_B T$) | $16.2 \pm 0.5$ | $14.5 \pm 0.7$ |
| *Simulations in the presence of ProTα* |  |  |
| $\tau_0$ (time steps) | $(35 \pm 12) \cdot 10^8$ | $(20 \pm 12) \cdot 10^8$ |
| $x^\ddagger$ (nm) | $0.51 \pm 0.03$ | $0.41 \pm 0.04$ |
| $\Delta G^\ddagger$ ($k_B T$) | $10.8 \pm 0.3$ | $9.6 \pm 0.3$ |

